# Discovery of antibiotics in the archaeome using deep learning

**DOI:** 10.1101/2024.11.15.623859

**Authors:** Marcelo D. T. Torres, Fangping Wan, Cesar de la Fuente-Nunez

**Author notes:** **Lead Contact:** Cesar de la Fuente-Nunez. These authors contributed equally (Marcelo D. T. Torres, Fangping Wan).

## Abstract

Antimicrobial resistance (AMR) is one of the greatest threats facing humanity, making the need for new antibiotics more critical than ever. While most antibiotics have traditionally been derived from bacteria and fungi, archaea—a distinct and underexplored domain of life—offer a largely untapped reservoir for antibiotic discovery. In this study, we leveraged deep learning to systematically explore the archaeome, uncovering promising new candidates for combating AMR. By mining 233 archaeal proteomes, we identified 12,623 molecules with potential antimicrobial activity. These newly discovered peptide compounds, termed archaeasins, exhibit unique compositional features that differentiate them from traditional antimicrobial peptides, including a distinct amino acid profile. We synthesized 80 archaeasins, 93% of which demonstrated antimicrobial activity *in vitro*. Notably, *in vivo* validation identified archaeasin-73 as a lead candidate, significantly reducing bacterial loads in mouse infection models, with effectiveness comparable to established antibiotics like polymyxin B. Our findings highlight the immense potential of archaea as a resource for developing next-generation antibiotics.

## Introduction

The rise of antimicrobial resistance (AMR) is one of the most urgent global health threats, as resistant pathogens undermine the efficacy of existing antibiotics, leading to increasingly difficult-to-treat infections. This growing crisis highlights the critical need for novel antibiotics^1^. However, the discovery pipeline for new antibiotics has slowed significantly in recent decades^1,2^, and has traditionally relied primarily on bacteria and fungi as sources. To address this challenge, there is a pressing need to explore alternative domains of life for potential antibiotic molecules.

Archaea, a distinct and ancient domain of life, have been largely overlooked in antibiotic discovery efforts. Despite their unique biochemical properties and resilience in extreme environments, archaea have not been systematically explored as a source of novel antibiotics. This represents a significant gap in our search for new antimicrobial agents, particularly in light of the diverse and unexplored nature of the archaeal domain.

Computational mining of biological information has been shown to be a promising approach for antibiotic discovery^3–7^. We previously mined the human proteome^4,7^ and extinct organisms^5,6^ as source of antibiotics and identified encrypted peptides (EPs), fragments within proteins that possess antimicrobial properties but do not necessarily are part of the immune system^7^. In this study, we used a deep learning approach to systematically explore the archaeome (**Fig. 1**) —the collective genetic material of archaea—as a novel and promising source of antibiotic molecules. We uncovered and characterized antibiotic candidates within this untapped reservoir, potentially offering new solutions to combat AMR. Our findings pave the way for the integration of archaea into the antibiotic discovery pipeline, presenting an underexplored source of potentially useful molecules.

**Figure 1.**
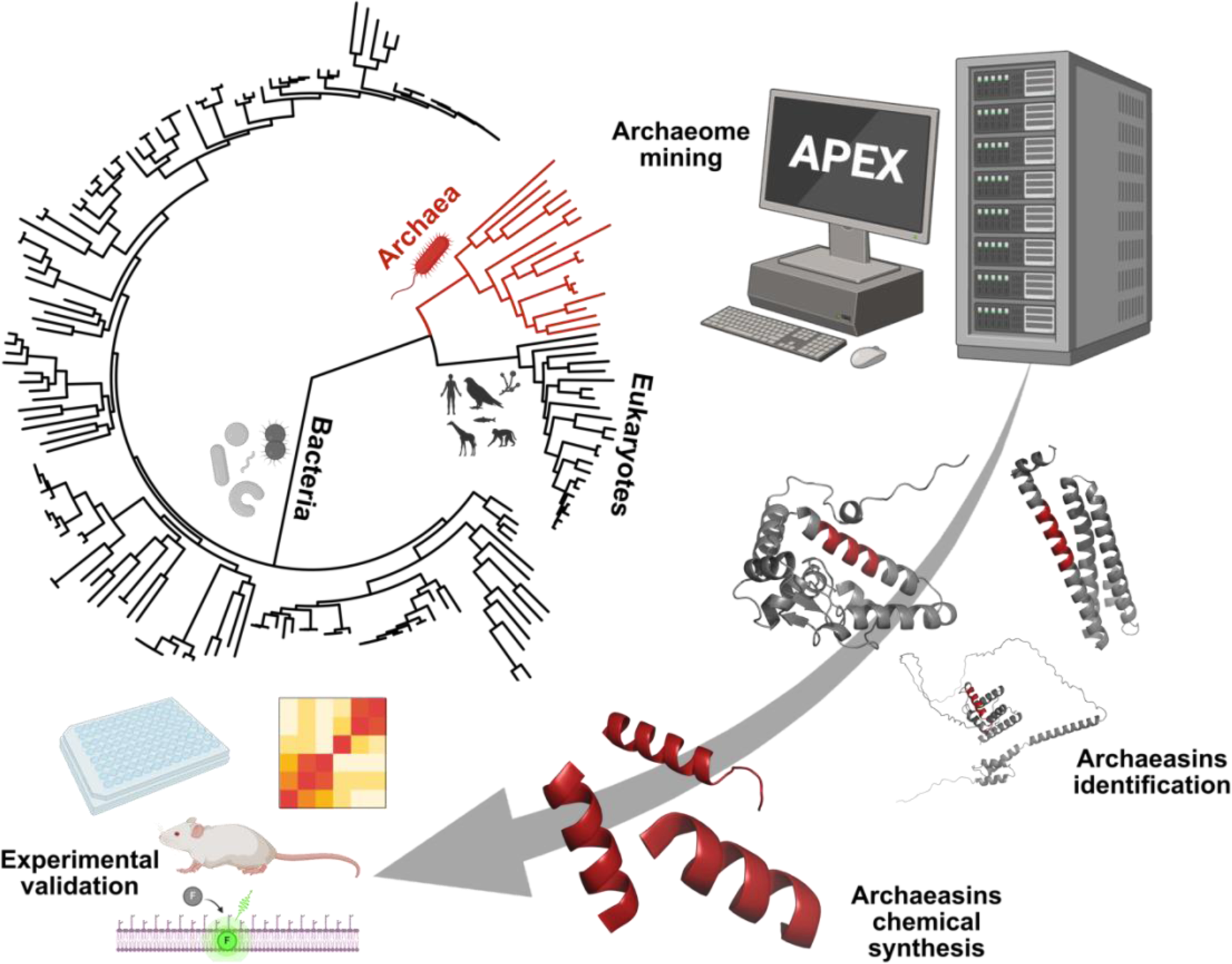
Discovery of antibiotics from Archaea using deep learning. The archaeome was systematically mined using our deep learning algorithm, APEX. Peptide sequences ranging from 8 to 50 amino acid residues within archaeal proteins were analyzed through multitask deep learning models trained on both public and in-house peptide datasets to predict antimicrobial activity. The top-ranked peptides, based on their predicted antimicrobial potential, were chemically synthesized and extensively evaluated against clinically relevant pathogens in both *in vitro* and animal model studies. Comprehensive assays were conducted to investigate the mechanism of action, toxicity, physicochemical properties, and potential synergistic interactions of these peptides. The protein and peptide structures depicted in the figure were created with PyMOL Molecular Graphics System, version 3.0 Schrödinger, LLC. Figure created with BioRender.com.

## Results

### Archaeasins identified by APEX 1.1

We collected 18,677 non-redundant protein sequences from 233 archaeal organisms available on UniProt^8^ and used APEX 1.1, a deep learning antimicrobial activity predictor^6^ retrained on updated data (see **APEX 1.1** in **Methods** section), to mine EPs within archaeal proteomes. As APEX predicted bacterial strain-specific minimum inhibitory concentrations (MICs), we used the mean MIC to represent the overall antimicrobial potency of the peptides and found 12,623 encrypted peptides with a mean MIC ≤100 μmol L^−1^ (**Fig. 2a** and **Data S1**).

**Figure 2.**
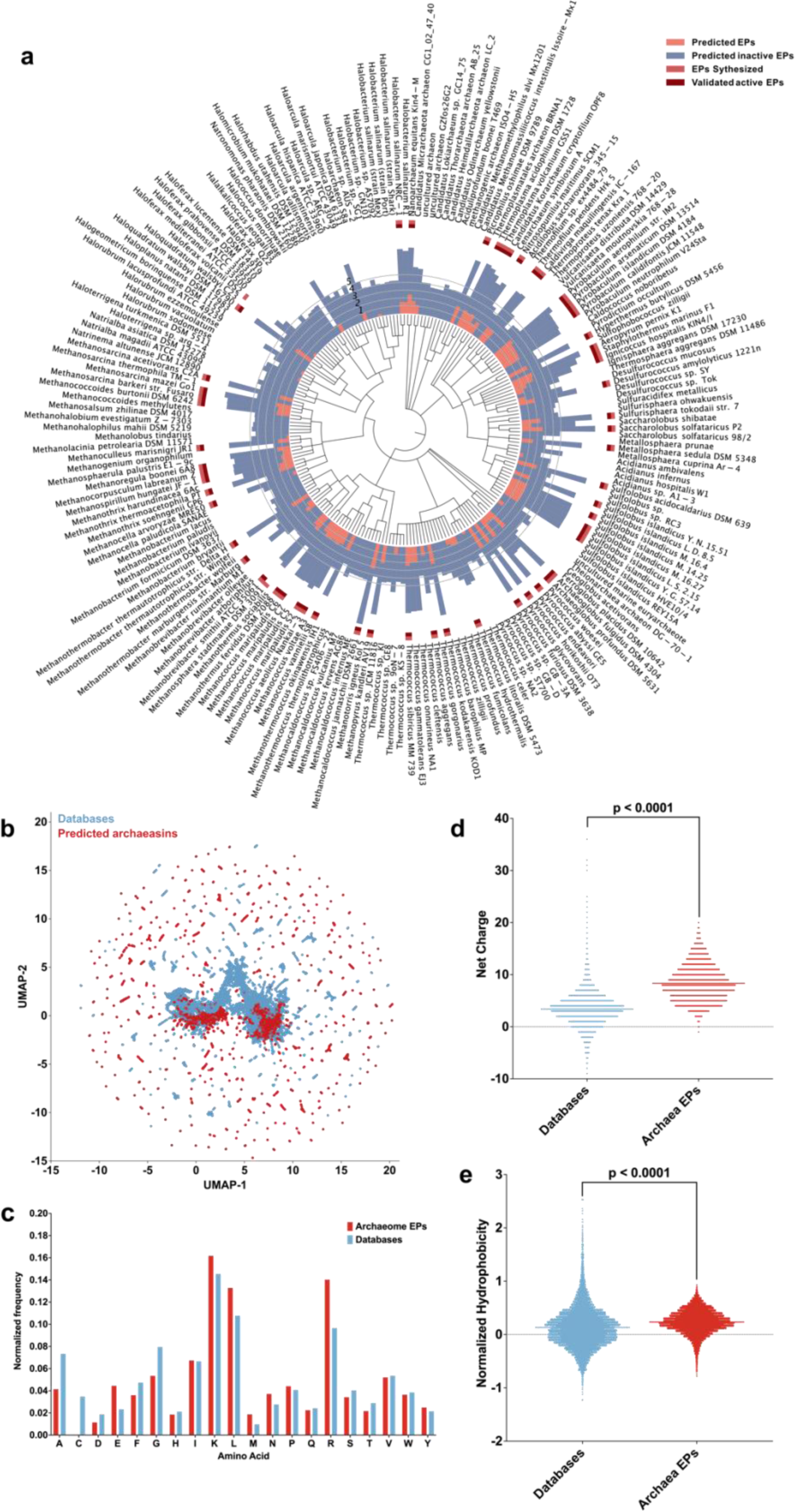
APEX exploration of the Archaeome. **(a)** Archaeal proteomes were systematically scanned to identify encrypted peptides with potential antimicrobial activity. Circular bars denote the log10-transformed average active (red) and inactive (blue) encrypted peptides discovered by APEX. A peptide was classified as active if its predicted mean MIC against tested bacterial strains was ≤100 μmol L^−1^. The values were normalized by the number of proteins per organism scanned. Organisms whose EPs were selected for further validation are highlighted in bold. Archaea with EPs that were synthesized are indicated by a light red square, and those experimentally validated as active are highlighted with a dark red square. **(b)** Sequence space exploration using a similarity matrix. The graph illustrates a bidimensional sequence space visualization of peptide sequences found in DBAASP and antimicrobial EPs discovered by APEX in Archaea organisms. Sequence alignment was used to generate a similarity matrix for all peptide sequences in DBAASP and the 12,623 antimicrobial EPs predicted by APEX (see also **Data S1** and **S2**). Each row in the matrix represents a feature representation of a peptide based on its amino acid composition. Uniform Manifold Approximation and Projection (UMAP) was applied to reduce the feature representation to two dimensions for visualization (see also **Fig. S1a**). **(c)** Comparison of amino acid frequency in Archaeal EPs with known antimicrobial peptides (AMPs) from the DBAASP, APD3, and DRAMP 3.0 databases (see also **Fig. S1b-e**). Distribution of two physicochemical properties for peptides with predicted antimicrobial activity, compared with AMPs from DBAASP, APD3, and DRAMP 3.0: net charge **(d)** and normalized hydrophobicity **(e)**. Net charge influences the initial electrostatic interactions between the peptide and negatively charged bacterial membranes, while hydrophobicity affects interactions with lipids in the membrane bilayers (see also **Fig. S2**). Statistical significance in **d** and **e** was determined using two-tailed t-tests followed by Mann–Whitney test; P values are shown in the graph. The solid line inside each box represents the mean value for each group.

To investigate the distribution of AMP-like encrypted peptides within the archaeal proteomes, we first performed sequence alignment on the combined set of 12,623 encrypted peptides and 19,775 publicly available AMPs DBAASP^9^, APD3^10^, and DRAMP^11^ (see **APEX 1.1** in **Methods** section). We then applied Uniform Manifold Approximation and Projection (UMAP)^12^ to reduce and visualize the sequence similarity matrix derived from the local sequence alignment (**Fig. 2b**).

The amino acid composition of archaeasins reveals distinctive features compared to known AMPs from databases (**Fig. 2c**) and other EPs from extinct organisms previously discovered by APEX (**Fig. S1b-e**). Notably, archaeasins contain an abundance of glutamic acid residues, surpassing levels typically found in known AMPs. This higher prevalence of negatively charged residues is also observed when comparing archaeasins to other extinct EPs. Despite this, archaeasins still maintain a prevalence of cationic residues, leading them to display a slightly higher proportion of cationic residues compared to database entries, suggesting a unique balance in charge distribution (**Fig. 2d**). Despite these differences, their hydrophobicity remains comparable to standard database sequences (**Fig. 2e**). Additionally, archaeasins show a tendency towards increased amphiphilicity, indicating a balanced distribution between hydrophobic and hydrophilic residues (**Fig. S2**). These compositional nuances distinguish archaeasins from other classes of peptide sequences studied.

### Antimicrobial activity assays

To experimentally validate the antimicrobial activity of the Archaea encrypted peptides, we selected 80 peptides that were both sequentially diverse and top-ranked by APEX 1.1 (**Data S2**). We prioritized peptides with less than <70% sequence similarity to known AMP sequences for chemical synthesis and experimental validation (**Fig. S1a**). Additionally, when two mined sequences exhibited high sequence similarity, we retained only the peptide with the higher predicted antimicrobial activity (see **Archaea encrypted peptides selection** in **Methods** section).

These archaeasins were tested against clinically relevant pathogens (*Acinetobacter baumannii*, *Escherichia coli*, *Klebsiella pneumoniae*, *Pseudomonas aeruginosa*, *Staphylococcus aureus, Enterococcus faecalis, and Enterococcus faecium*) at a range of concentrations from 1 to 64 μmol L⁻¹. The results showed that 75 out of the 80 encrypted peptides exhibited antimicrobial activity (MIC ≤64 μmol L^−1^) against at least one pathogenic strain (**Fig. 3a**), resulting in a hit rate of over 93%. Additionally, the Pearson correlation (ρ = 0.503) between predicted and experimentally validated MICs demonstrated the predictive power of APEX 1.1 (**Fig. S3**). When comparing the Pearson and Spearman correlations between experimental and predicted MICs from the first version of APEX^6^ to APEX 1.1, used to explore the archaeome, on the 80 archaeasins synthesized, we observed that APEX 1.1 significantly outperformed APEX (**Supplementary Tables S1** and **S2**).

**Figure 3.**
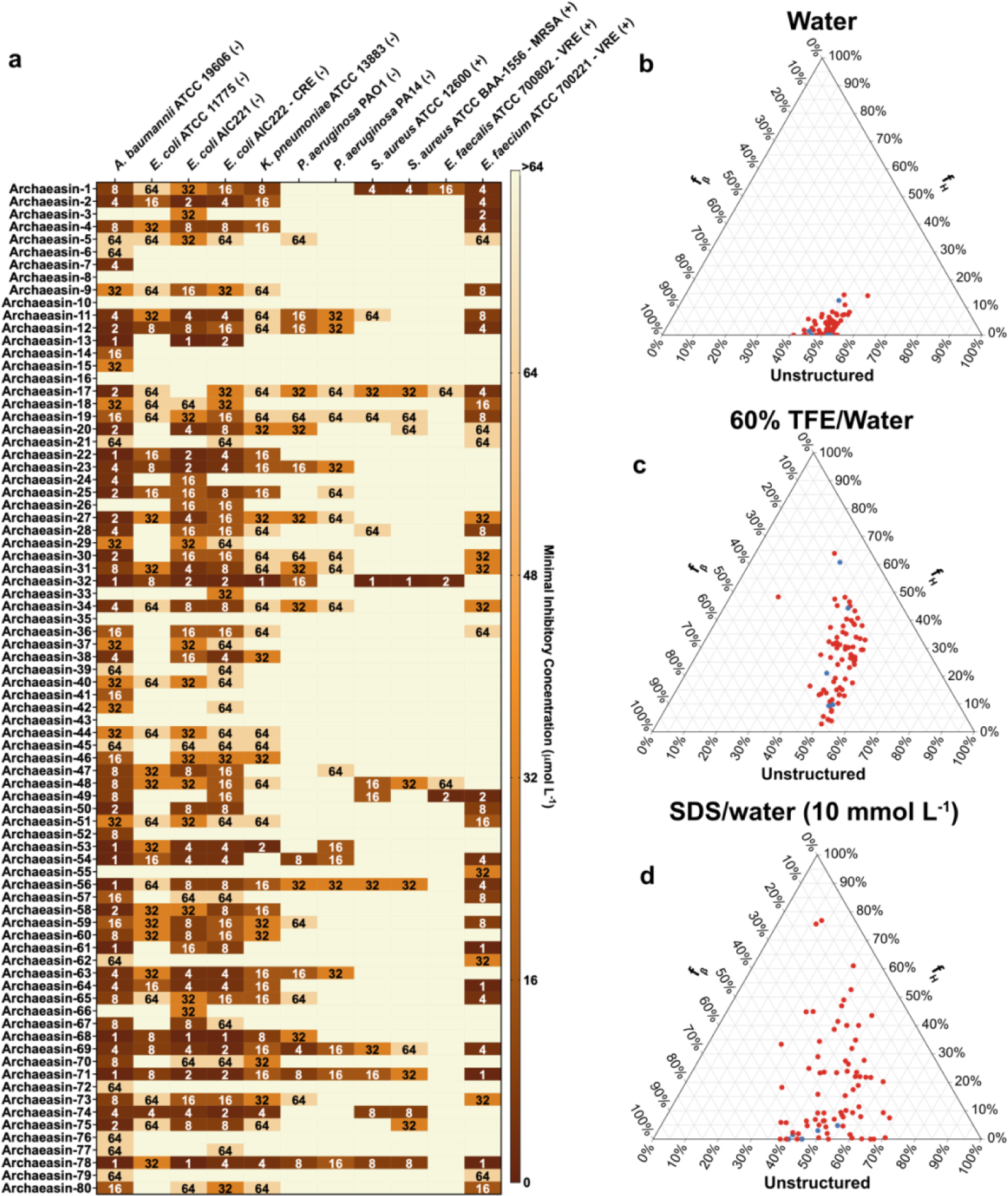
Antimicrobial activity and secondary structure profiles of antibiotics from the archaeome. **(a)** Heat map displaying the antimicrobial activities (μmol L^−1^) of active antimicrobial agents from Archaea against 11 clinically relevant pathogens, including antibiotic-resistant strains. Briefly, 10^5^ bacterial cells were incubated with serially diluted EPs (0–64 μmol L^−1^) at 37 °C. Bacterial growth was assessed by measuring the optical density at 600 nm in a microplate reader one day post-treatment. The MIC values presented in the heat map represent the mode of the replicates for each condition (see also **Fig. S3**). **(b)** Ternary plots showing the percentage of secondary structure for each peptide (at 50 μmol L^−1^) in three different solvents: water, 60% trifluoroethanol (TFE) in water, and Sodium dodecyl sulfate (SDS, 10 mmol L^−1^) in water. Secondary structure fractions were calculated using the BeStSel server^16^. Red dots indicate active archaeasins, while blue dots represent inactive peptides (see also **Fig. S4**).

### Secondary structure studies

The secondary structure of short peptides is often dynamic, transitioning between disordered and ordered conformations at hydrophobic/hydrophilic interfaces. These structural transitions are critical in determining the antimicrobial and other biological functions of peptides. To assess the secondary structure of the synthesized archaeasins, we conducted Circular Dichroism experiments in various environments: water, sodium dodecyl sulfate (SDS)/water (10 mmol L^−1^), and trifluoroethanol (TFE)/water (3:2, v:v), mixture. SDS micelles were chosen as a membrane-mimetic environment because of the lipid bilayers environment that is similar to biological bilayers^13^. The TFE/water mixture is known to induce α-helical structures by dehydrating the amide groups in the peptide backbone, thus favoring intramolecular hydrogen bonds that promote a helical conformation^14,15^. All archaeasins were tested at 50 μmol L^−1^ in the wavelength range of 260 to 190 nm (**Fig. S4**). To determine secondary conformation fractions, we used the Beta Structure Selection (BeStSel) server^16^ (**Fig. 3b-d**). As expected, given that archaeasins are short sequences (<50 amino acid residues), all tested peptides were unstructured in water (**Fig. 3b**), with a slight tendency toward β-like structures (20% < f_β_ < 45%) in the other two analyzed media, the helical-inducer trifluoroethanol and water mixture (3:2, v:v; **Fig. 3c**) and sodium dodecyl sulfate (SDS) micelles (10 mmol L^−1^) in water (**Fig. 3d**). This behavior is typical for short peptides exhibiting antimicrobial activity^17–19^. While encrypted peptides primarily adopt β-like structures^3,20^, archaeasins displayed helical conformations in helical-inducing media and upon interaction with lipid bilayers.

### Synergy assays

To explore whether molecules from the same archaeal strains or their closest relatives could synergize and potentiate each other’s antimicrobial activity against pathogens, we performed checkerboard assays. These assays tested peptide concentrations ranging from twice the MIC to concentrations up to 32-times lower, under the same conditions as those used for the antimicrobial assays. We initially selected the bacterial strain *A. baumannii* ATCC 19606, known for its high antibiotic resistance and significant role as an opportunistic nosocomial pathogen with substantial global mortality rates^21^. This strain was particularly susceptible to the archaeasins. We then selected peptides from strains closely related on the phylogenetic tree (pairwise distance ≤8), resulting in the testing of 79 pairs of archaeasins (**Fig. 4**).

**Figure 4.**
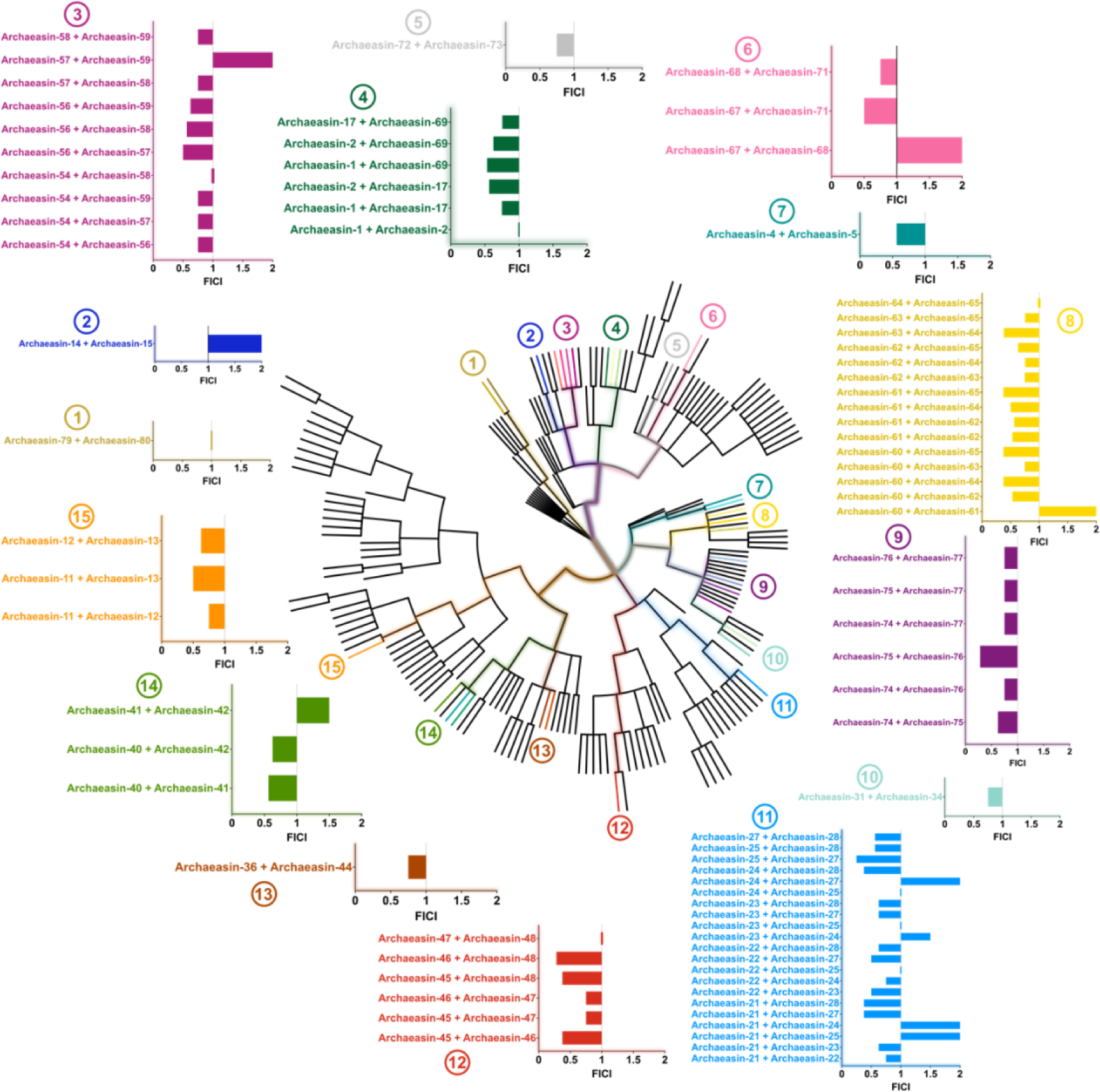
Synergistic interaction between Archaeal peptide antibiotics. The synergistic interactions between pairs of EPs from the same or closely related organisms (phylogenetic pairwise distance ≤8) that exhibited activity against *A. baumannii* ATCC 19606 were assessed using checkerboard assays. These assays involved 2-fold serial dilutions, ranging from 2×MIC to MIC/32. The histogram displays the fractional inhibitory concentration index (FICI) values obtained for each pair of EPs. A total of 79 pairs were evaluated. Low FICI values (≤0.5) indicate synergistic interactions, intermediary values (0.5<FICI≤1) indicate additive effects, and higher values (1<FICI≤2) indicate indifferent interactions.

Most of the combinations tested exhibited synergistic or additive interactions, as determined by the fractional inhibitory concentration index^22^ (FICI). Notably, archaeasins from *Methanocaldococcus* species demonstrated some of the lowest FICI values, ranging from 0.25 (archaeasins 25 and 27) to 0.375 (archaeasins 21 and 27, 21 and 28, and 24 and 28). Similarly, *Methanothermobacter* species compounds showed FICI values from 0.28 (archaeasins 46 and 48) to 0.375 (archaeasins 45 and 46, 46 and 48). *Thermococcus* species had a FICI of 0.28 (archaeasins 75 and 76), while compounds derived from *Pyrococcus* species displayed a FICI of 0.375 for combinations of archaeasins 60 and 64, 60 and 65, 61 and 65, and 63 and 64.

### Mechanism of action studies

To understand how archaeasins exert their effect on bacterial cells, we conducted fluorescence assays to determine if their mechanism of action involves membrane targeting. First, we identified 70 antimicrobial hits among archaeasins effective against *A. baumannii* ATCC 19606 (**Fig. 3a**). We then assessed the ability of these peptides, at their MIC values, to permeabilize (**Fig. 5a and S5a**) and depolarize (**Fig. 5b and S5b**) bacterial outer and cytoplasmic membranes, respectively.

**Figure 5.**
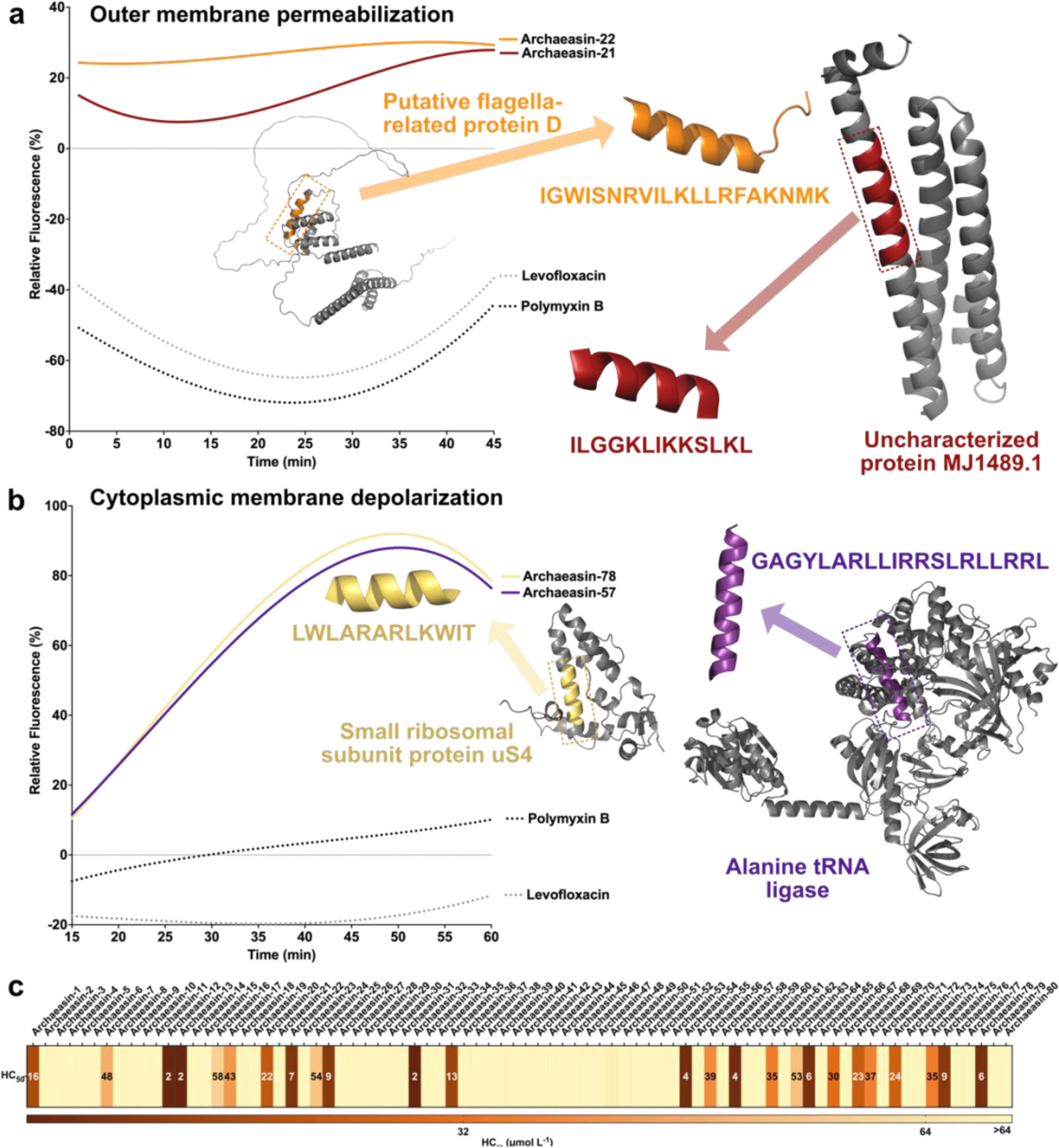
Mechanism of action and hemolytic activity of antibiotics from the archaeome. To assess whether Archaea EPs act on bacterial membranes, all active peptides against *A. baumannii* ATCC 19606 were subjected to outer membrane permeabilization and cytoplasmic membrane depolarization assays. The fluorescent probe 1-(N-phenylamino)naphthalene (NPN) was used to assess membrane permeabilization **(a)** induced by the tested EPs (see also **Fig. S5a**). The fluorescent probe 3,3′-dipropylthiadicarbocyanine iodide (DiSC3-5) was used to evaluate membrane depolarization **(b)** caused by archaeasins (see also **Fig. S5b**). The values displayed represent the relative fluorescence of both probes, with non-linear fitting compared to the baseline of the untreated control (buffer + bacteria + fluorescent dye) and benchmarked against the antibiotics polymyxin B and levofloxacin. **(c)** Hemolytic concentrations leading to 50% cell lysis (HC50) were determined by interpolating the dose-response data using a non-linear regression curve. All experiments were performed in three independent replicates (see also **Fig. S6**). The protein and peptide structures depicted in the figure were created with PyMOL Molecular Graphics System, version 3.0 Schrödinger, LLC.

To evaluate the ability of archaeasins to permeabilize the outer membrane of Gram-negative bacteria, we employed 1-(N-phenylamino)naphthalene (NPN) assays. NPN is a lipophilic dye that fluoresces in lipid-rich environments, such as bacterial outer membranes. Damage to the bacterial outer membrane allows NPN to penetrate, increasing fluorescence (**Fig. 5a**). Only archaeasins 21 (parent protein: uncharacterized protein MJ1489.1) and 22 (parent protein: putative flagella-related protein D) from *Methanocaldococcus jannaschii* effectively permeabilized the bacterial outer membrane. Polymyxin B served as a positive control in these experiments^4^. Overall, archaeasins did not permeabilize the bacterial outer membrane to the extent observed for AMPs^23,24^ or other human- or animal-derived EPs^4,6^.

We then used 3,3′-dipropylthiadicarbocyanine iodide (DiSC3-5), a fluorophore that indicates cytoplasmic membrane depolarization. Disruption of the transmembrane potential causes the fluorophore to migrate to the extracellular space, resulting in increased fluorescence. Among the 70 peptides tested, 34 archaeasins significantly depolarized the cytoplasmic membrane more than the control group treated with polymyxin B^4^ (**Fig. 5b**). Archaeasins 78 (parent protein: alanine tRNA ligase) from *Thermofilum pendens* and 57 (parent protein: small ribosomal subunit protein uS4) from *Pyrobaculum arsenaticum* were particularly effective depolarizers.

These findings suggest that archaeasins primarily exert their antimicrobial effects by depolarizing the cytoplasmic membrane, rather than permeabilizing the outer membrane. This suggests a mechanism akin to that of the recently reported SEPs^20^ and unusual for conventional AMPs ^23,24^ and EPs^4^, which typically target the outer membrane^17^.

### Hemolytic activity assays

To assess the potential toxicity of the synthesized archaeasins, we exposed them to human red blood cells (RBCs), a common method for evaluating the toxicity of antimicrobial agents^18,24,25^. Twenty five out of the 80 (31.3%) archaeasins tested showed moderate to low hemolytic activity within the explored concentration range, *i.e.*, their HC50 values (linear regression of the peptide concentration that leads to 50% RBCs lysis) were ≤64 μmol L^−1^ (**Figs. 5c and S6**). Most sequences active against bacterial pathogens at low MIC values did not display toxic effects at those concentrations (**Fig. S6**). However, seven archaeasins, specifically archaeasins 12, 13, 32, 54, 58, 64 and 78, did show toxic effects.

### Anti-infective activity of encrypted peptides in preclinical animal models

To evaluate whether the lead archaeasins retained their antimicrobial potency in complex living systems, we tested them in two mouse models: a skin abscess^26–28^ and a deep thigh infection model^4,5^ (**Figure 6a**). In both models, we used *A. baumannii*, a pathogen responsible for infections in the blood, urinary tract, lungs, and topical wounds, and a major cause of mortality in hospitalized patients due to its antimicrobial resistance^29^. Three lead archaeasins displayed potent activity against *A. baumannii* and no cytotoxicity (CC50 <64 μmol L^−1^): archaeasin-2 (MIC value = 4 μmol L^−1^) from *Aeropyrum pernix*, archaeasin-17 (MIC value = 2 μmol L^−1^) from *Ignicoccus hospitalis*, and archaeasin-73 (MIC value = 8 μmol L^−1^) from *Sulfurisphaera tokodaii*.

**Figure 6.**
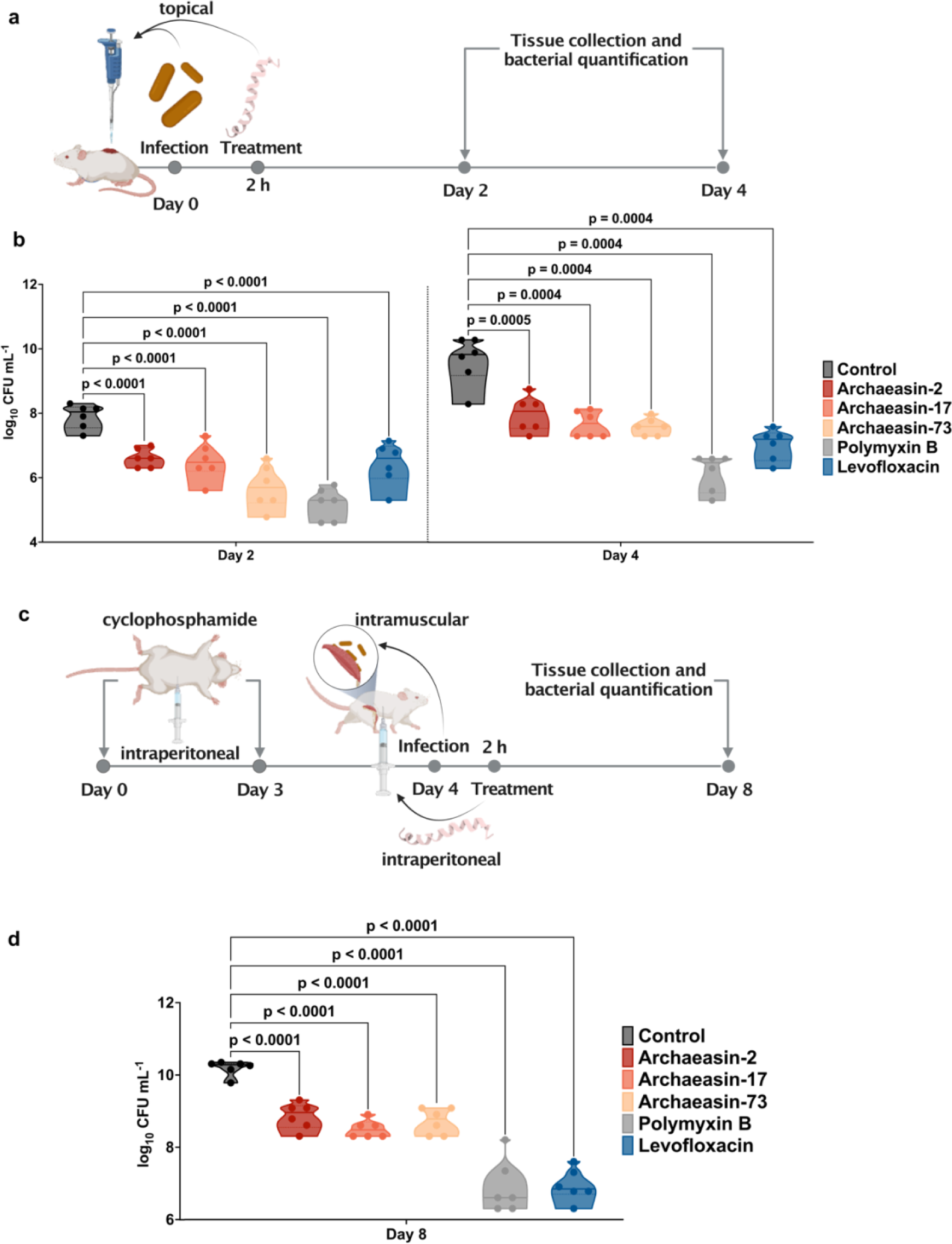
Anti-infective activity of archaeasins in animal models. **(a)** Schematic representaton of the skin abscess mouse model used to assess the anti-infective activity of archaeasins (n = 6) against *A. baumannii* ATCC 19606. **(b)** Archaeasin-2, archaeasin-17, and archaeasin-73, administered at their MIC in a single dose post-infection, inhibited the proliferation of the infection for up to 4 days after treatment compared to the untreated control group. Notably, archaeasin-73 reduced the infection in some mice, demonstrating activity comparable to the control antibiotic, polymyxin B. **(c)** Schematic of the neutropenic thigh infection mouse model, where archaeasins were administered intraperitoneally. Anti-infective activity against *A. baumannii* ATCC 19606 was evaluated 4 days after intraperitoneal peptide administration (n = 6). **(d)** Four days after intraperitoneal injection, all archaeasins at their MIC exhibited a bacteriostatic effect, containing the *A. baumannii* ATCC 19606 infection, though their activity was less potent than that of polymyxin B and levofloxacin, compared to the untreated control group (see also **Figure S7**). Statistical significance in panels **b** and **d** was determined using one-way ANOVA followed by Dunnett’s test; P values are shown in the graphs. In the violin, the center line represents the mean, the box limits the first and third quartiles, and the whiskers (minima and maxima) represent 1.5 × the interquartile range. The solid line inside each box represents the mean value obtained for each group. Panels **a** and **c** were created with BioRender.com.

In the skin abscess model, infection was established with a 20 μL bacterial load of 1.2×10^5^ of *A. baumannii* cells in phosphate buffer solution (PBS) applied to a wounded area of the skin (**Figure 6a**). A single dose of each archaeasin at their respective MIC was administered to the infected area. Two days post-infection, all archaeasins tested showed significantly reduced bacterial counts by 1.5 to 2 orders of magnitude. Archaeasin-73, in particular, reduced the bacterial load by two orders of magnitude compared to the untreated control group. Its potency was comparable to that observed in the positive control group of mice treated with polymyxin B and was higher than that of the levofloxacin control group (**Figure 6b**). Four-days post infection, all archaeasins and the two antibiotics, polymyxin B and levofloxacin, continued to prevent bacterial growth with similar efficacy. Polymyxin B reduced bacterial counts by four orders of magnitude compared to the untreated control group of mice, while all other treatment groups showed a two- to three-order magnitude decrease. These results are promising, as the archaeasins were administered only once after the abscess had been established, highlighting their anti-infective potential. Importantly, no significant changes in weight, used as a proxy for toxicity, were observed in our experiments (**Figure S7a**).

Next, we assessed the efficacy of the same lead archaeasins (archaeasin-2, archaeasin-17, and archaeasin-73) in a murine deep thigh infection model (**Figure 6c**), which is widely used to assess the antibiotic potential of compounds. Mice were administered two rounds of cyclophosphamide treatment for immunosuppression before the intramuscular infection with 1×10^5^ cells in 100 μL of *A. baumannii*. A single dose of each archaeasin (at their MIC) was delivered intraperitoneally (**Figure 6c**). Four days post-treatment, the archaeasins were unable to prevent the growth of the infection, while the antibiotics polymyxin B and levofloxacin (positive controls) reduced the bacterial load by three orders of magnitude (**Figure 6d**). Four days post-treatment, the bacterial counts remained stable for all peptide treatment conditions and the treatments with polymyxin-B and levofloxacin, while the untreated control increased by two orders of magnitude. No significant changes in weight were observed, indicating that the archaeasins are non-toxic (**Figure S7b**). These *in vivo* results support the antibiotic properties of archaeasins under physiological conditions and provide a strong foundation for advancing their development as potential antimicrobial agents.

## Discussion

In this study, we systematically explored the archaeome using the deep learning model APEX 1.1, revealing a wealth of previously unrecognized antibiotic molecules within Archaea. Our findings highlight the untapped potential of Archaea as a source of novel antimicrobial agents^30–36^, expanding the traditional focus beyond bacteria and fungi, which have historically been the primary sources of antibiotics derived from nature.

We report the discovery of archaeasins, a new class of antimicrobial peptides with unique compositional characteristics. The distinctive amino acid profiles of these peptides, including an unusually high abundance of glutamic acid residues and a balanced distribution of cationic and amphiphilic residues, suggest that archaeasins may represent a new paradigm in peptide-based antibiotic design.

Our synergy assays further underscored the potential of archaeasins to work in concert, enhancing their antimicrobial efficacy when combined. The low FICI values observed in combinations from closely related strains, particularly within the *Methanocaldococcus* and *Methanothermobacter* species, point to the possibility of developing combination therapies that leverage these synergistic effects. Such combinations could provide more effective treatment options, especially against multidrug-resistant pathogens.

Mechanism of action studies revealed that archaeasins primarily exert their antimicrobial effects by depolarizing the bacterial cytoplasmic membrane, rather than by permeabilizing the outer membrane. This finding is particularly intriguing, as it suggests that archaeasins may operate through a mechanism distinct from that of conventional AMPs, which often target the outer membrane. Depolarization of the cytoplasmic membrane is a critical process that disrupts bacterial homeostasis, leading to cell death. Interestingly, this mode of action aligns more closely with that of recently described small open reading frame-encoded peptides (SEPs)^20^.

Our *in vivo* experiments in mouse models demonstrated that archaeasins retain their antimicrobial potency in complex biological systems, effectively reducing bacterial loads in both skin abscess and deep thigh infection models. The observed efficacy, particularly with archaeasin-73, which showed results comparable to than traditional antibiotics like polymyxin B and levofloxacin, is promising. These results suggest that archaeasins have the potential to be developed into viable therapeutic agents, especially for infections caused by multidrug-resistant pathogens such as *A. baumannii*. Importantly, the lack of significant toxicity observed in these models further supports the safety profile of these peptides, a crucial consideration for future development.

In conclusion, our study demonstrates the promise of using deep learning to unlock the archaeome as a source of novel antibiotics. The discovery of archaeasins warrants further development of these agents, opening a new frontier in the fight against antimicrobial resistance.

### Limitations of the study

Despite the promising results, several challenges and limitations remain. For instance, while the *in vivo* findings are encouraging, further studies are necessary to systematically evaluate the long-term efficacy and safety of archaeasins, including their pharmacokinetics, pharmacodynamics, and potential immunogenicity in humans.

Additionally, there are inherent limitations in using deep learning to explore archaeal proteomes as a framework for antibiotic discovery. Our current deep learning model is sequence-based and lacks structural information. While this approach allows for rapid analysis across proteomes, incorporating structural and three-dimensional descriptors in future iterations could improve the model’s accuracy in predicting antimicrobial activity. Another challenge is the limited availability of information on archaeal proteins. The virtual screening of Archaea proteomes was performed only on the high quality reviewed sequences from UniProt, while the unreviewed sequences were not included in the analysis. This inevitably led to an imbalanced analysis, with some interesting clades (e.g., DPANN and Asgardarchaeota) not being well characterized by APEX. In the future, we will apply APEX to screen unreviewed Archaea proteins to complement our current study.

Despite these limitations, this study serves as a proof of concept, demonstrating the potential of combining deep learning with extensive wet-lab validation, both *in vitro* and in animal models, to explore the archaeome for antimicrobial molecules. Our approach highlights an underexplored source of antibiotics and suggests that the integration of artificial intelligence with the study of archaeal organisms could lead to other significant medicinal discoveries.

## Acknowledgments

Cesar de la Fuente-Nunez holds a Presidential Professorship at the University of Pennsylvania and acknowledges funding from the Procter & Gamble Company, United Therapeutics, a BBRF Young Investigator Grant, the Nemirovsky Prize, Penn Health-Tech Accelerator Award, and the Dean’s Innovation Fund from the Perelman School of Medicine at the University of Pennsylvania. Research reported in this publication was supported by the Langer Prize (AIChE Foundation), the National Institute of General Medical Sciences of the National Institutes of Health under award number R35GM138201, and the Defense Threat Reduction Agency (DTRA; HDTRA11810041, HDTRA1-21-1-0014, and HDTRA1-23-1-0001). We thank Dr. Mark Goulian for kindly donating the following strains: *Escherichia coli* AIC221 [*Escherichia coli* MG1655 phnE_2::FRT (control strain for AIC222)] and *Escherichia coli* AIC222 [*Escherichia coli* MG1655 pmrA53 phnE_2::FRT (polymyxin resistant)]. We thank de la Fuente Lab members for insightful discussions. Figures created with BioRender.com are attributed as such. Molecules were rendered using the PyMOL Molecular Graphics System, Version 3.0 Schrödinger, LLC.

## Author contributions

Conceptualization: MDTT, FW, CFN

Methodology: MDTT, FW, CFN

Experimental investigation: MDTT

Computational investigation: FW

Visualization: MDTT, FW

Funding acquisition: CFN Supervision: CFN

Formal analysis: MDTT, FW

Writing – original draft: MDTT, FW, CFN

Writing – review & editing: MDTT, FW, CFN

## STAR Methods

### Key resources table

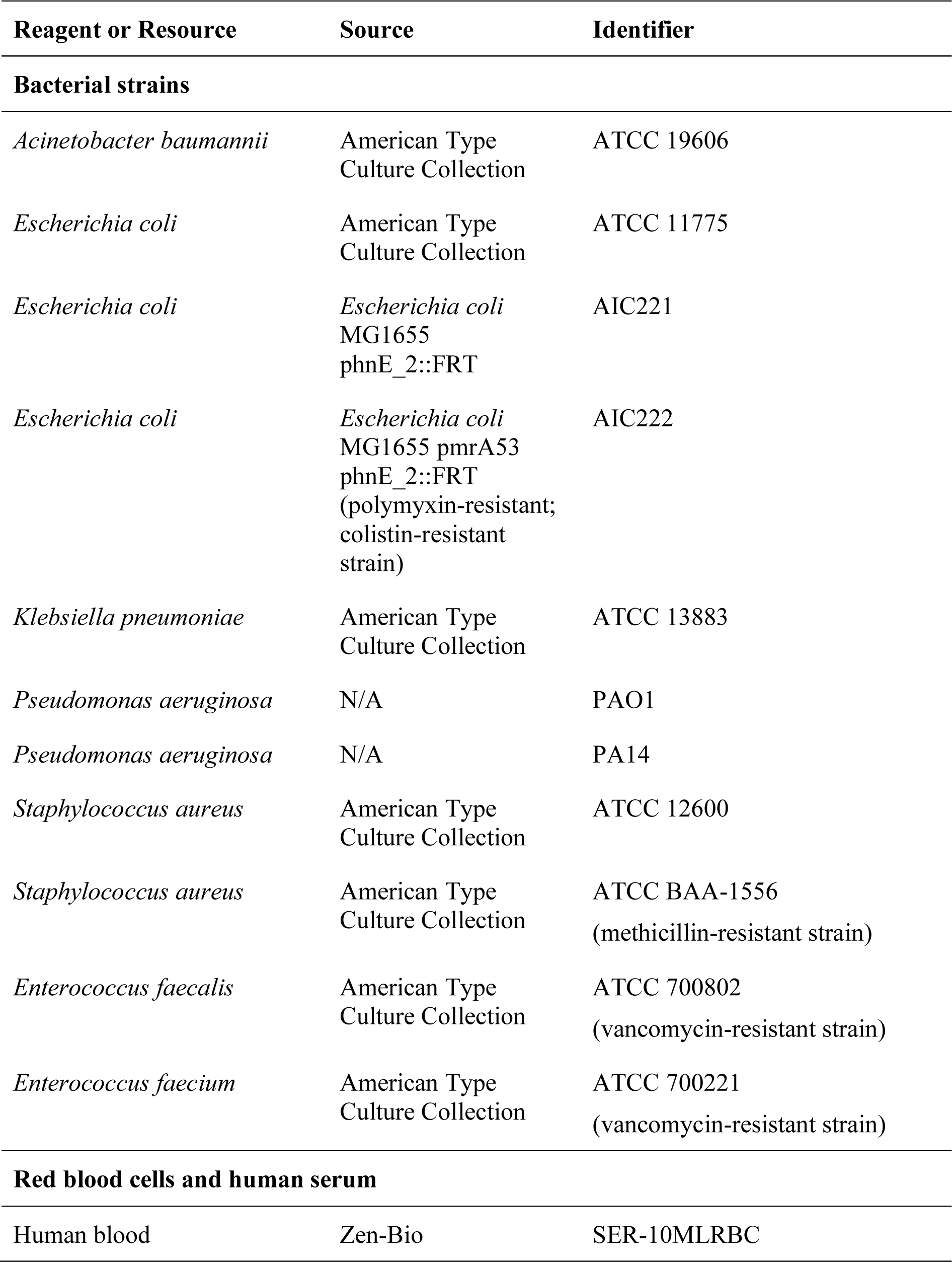
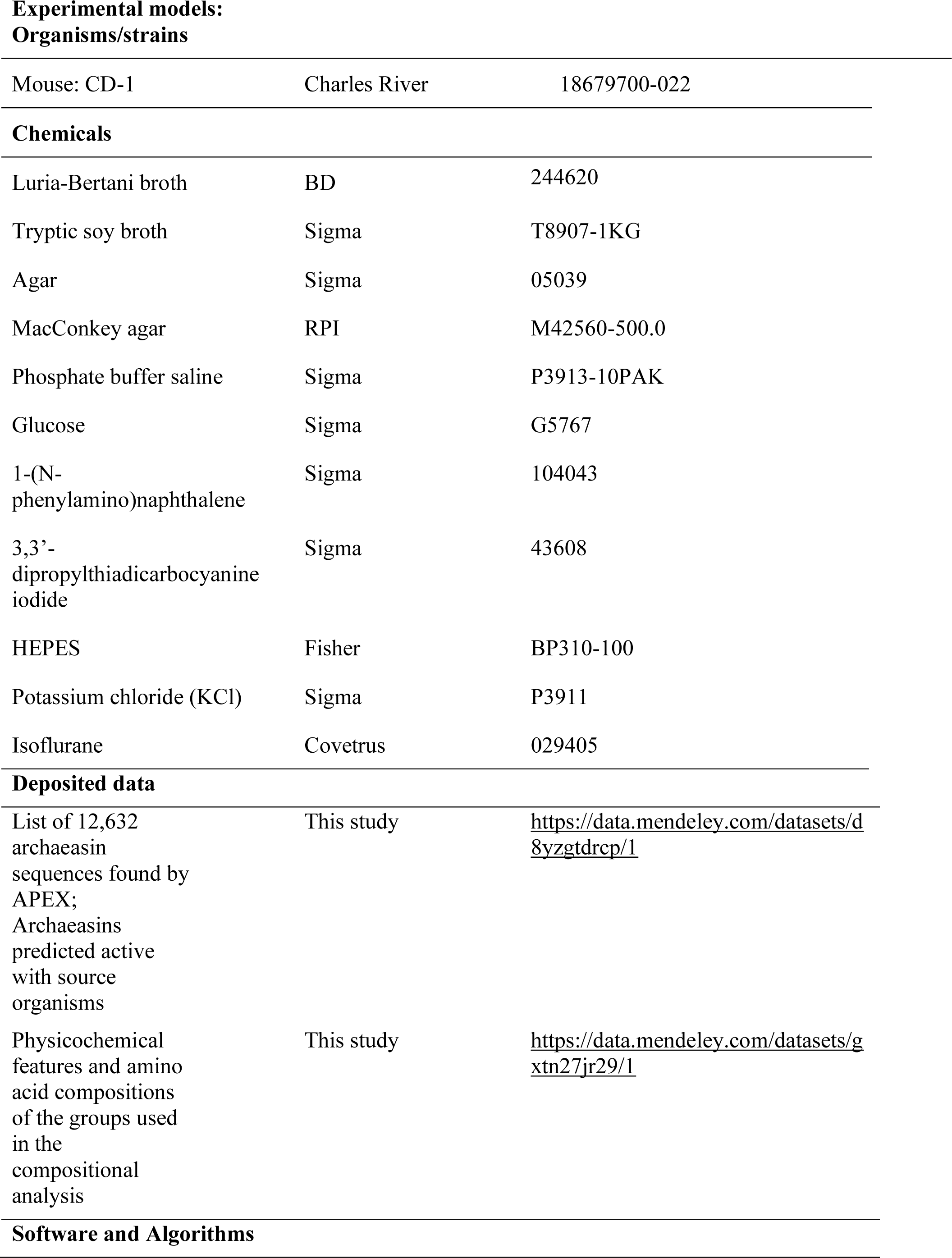
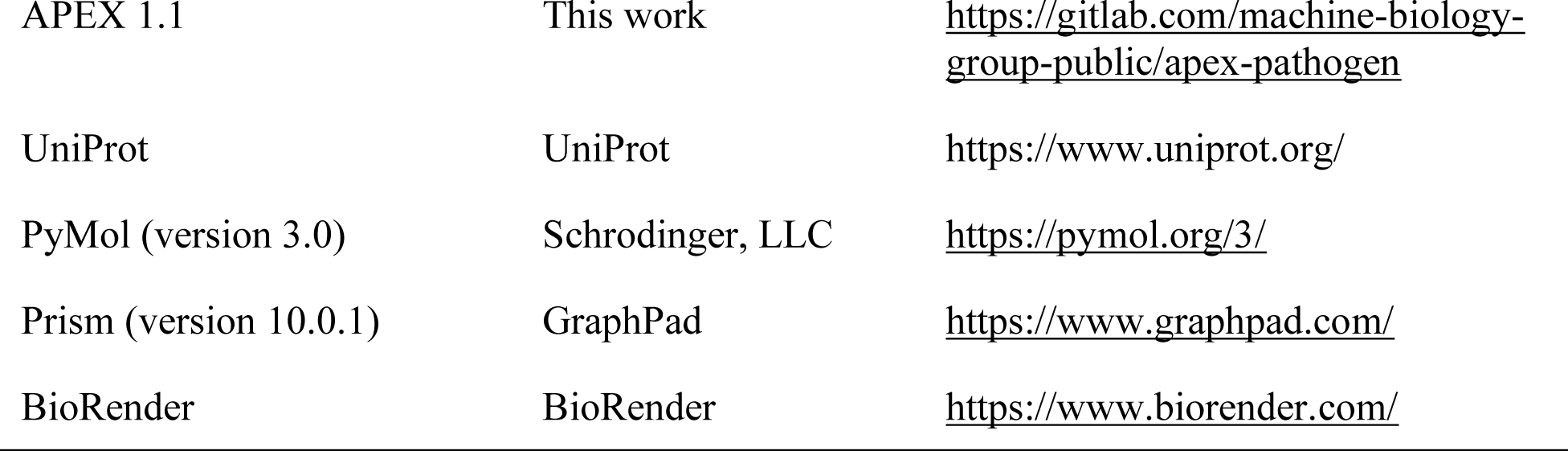

| Reagent or Resource | Source | Identifier |
| --- | --- | --- |
| **Bacterial strains** |  |  |
| *Acinetobacter baumannii* | American Type Culture Collection | ATCC 19606 |
| *Escherichia coli* | American Type Culture Collection | ATCC 11775 |
| *Escherichia coli* | *Escherichia coli* MG1655 phnE_2::FRT | AIC221 |
| *Escherichia coli* | *Escherichia coli* MG1655 pmrA53 phnE_2::FRT (polymyxin-resistant; colistin-resistant strain) | AIC222 |
| *Klebsiella pneumoniae* | American Type Culture Collection | ATCC 13883 |
| *Pseudomonas aeruginosa* | N/A | PAO1 |
| *Pseudomonas aeruginosa* | N/A | PA14 |
| *Staphylococcus aureus* | American Type Culture Collection | ATCC 12600 |
| *Staphylococcus aureus* | American Type Culture Collection | ATCC BAA-1556 (methicillin-resistant strain) |
| *Enterococcus faecalis* | American Type Culture Collection | ATCC 700802 (vancomycin-resistant strain) |
| *Enterococcus faecium* | American Type Culture Collection | ATCC 700221 (vancomycin-resistant strain) |
| **Red blood cells and human serum** |  |  |
| Human blood | Zen-Bio | SER-10MLRBC |
| **Experimental models: Organisms/strains** |  |  |
| Mouse: CD-1 | Charles River | 18679700-022 |
| **Chemicals** |  |  |
| Luria-Bertani broth | BD | 244620 |
| Tryptic soy broth | Sigma | T8907-1KG |
| Agar | Sigma | 05039 |
| MacConkey agar | RPI | M42560-500.0 |
| Phosphate buffer saline | Sigma | P3913-10PAK |
| Glucose | Sigma | G5767 |
| 1-(N-phenylamino)naphthalene | Sigma | 104043 |
| 3,3'-dipropylthiadicarbocyanine iodide | Sigma | 43608 |
| HEPES | Fisher | BP310-100 |
| Potassium chloride (KCl) | Sigma | P3911 |
| Isoflurane | Covetrus | 029405 |
| **Deposited data** |  |  |
| List of 12,632 archaeasin sequences found by APEX; Archaeasins predicted active with source organisms | This study | https://data.mendeley.com/datasets/d8yzgtdrcp/1 |
| Physicochemical features and amino acid compositions of the groups used in the compositional analysis | This study | https://data.mendeley.com/datasets/gxtn27jr29/1 |
| **Software and Algorithms** |  |  |
| APEX 1.1 | This work | https://gitlab.com/machine-biology-group-public/apex-pathogen |
| UniProt | UniProt | https://www.uniprot.org/ |
| PyMol (version 3.0) | Schrodinger, LLC | https://pymol.org/3/ |
| Prism (version 10.0.1) | GraphPad | https://www.graphpad.com/ |
| BioRender | BioRender | https://www.biorender.com/ |

### Resource Availability

#### Lead contact

Further information and requests for resources should be directed to and will be fulfilled upon reasonable request by the lead contact, Cesar de la Fuente-Nunez.

#### Materials availability

This study did not generate new unique reagents.

#### Data and code availability

- All reviewed canonical and isoform sequences of Archaea (Taxon ID: 2157) can be downloaded from UniProt (https://www.uniprot.org/).
- All original code and DOIs are publicly available and listed in the key resources table. APEX 1.1 is available at GitLab (https://gitlab.com/machine-biology-group-public/apex-pathogen) and Data S1 and Data S2 files are publicly available in Mendeley Data (https://data.mendeley.com/datasets/d8yzgtdrcp/1) and listed in the key resources table.
- Any additional information required to reanalyze the data reported in this paper is available from the lead contact upon reasonable request.

## Methodology

### Bacterial strains and growth conditions

In this study, we used the following pathogenic bacterial strains: *Acinetobacter baumannii* ATCC 19606, *Escherichia coli* AIC221 [*Escherichia coli* MG1655 phnE_2::FRT (control strain for AIC 222)] and *Escherichia coli* AIC222 [*Escherichia coli* MG1655 pmrA53 phnE_2::FRT (polymyxin resistant; colistin-resistant strain)], *Klebsiella pneumoniae* ATCC 13883, *Pseudomonas aeruginosa* PAO1, *Pseudomonas aeruginosa* PA14, and *Staphylococcus aureus* ATCC 12600. Pseudomonas Isolation (*Pseudomonas aeruginosa* strains) agar plates were exclusively used in the case of *Pseudomonas* species. All the other pathogens were grown in Luria-Bertani (LB) broth and on LB agar. In all the experiments, bacteria were inoculated from one-isolated colony and grown overnight (16 h) in liquid medium at 37 °C. In the following day, inoculums were diluted 1:100 in fresh media and incubated at 37 °C to mid-logarithmic phase.

### Red blood cells and human serum

Red blood cells (RBCs) and human serum were purchased from Zen-Bio. The RBC samples were obtained from the same certified healthy donor (blood type A^−^).

### Skin abscess infection mouse model

The back of six-week-old female CD-1 mice under anesthesia were shaved and injured with a superficial linear skin abrasion made with a needle. An aliquot of *A. baumannii* ATCC 19606 (6.3×10^5^ CFU mL^−1^; 20 μL) previously grown in LB medium until 0.5 OD mL^−1^ (optical value at 600 nm) and then washed twice with sterile PBS (pH 7.4, 13,000 rpm for 3 min) was added to the scratched area. Peptides diluted in sterile water at their MIC value were administered to the wounded area 1 h post-infection. Two- and four-days post-infection, animals were euthanized, and a uniform excision of the scarified skin was excised, homogenized using a bead beater (25 Hz for 20 min), 10-fold serially diluted, and plated on McConkey agar plates for CFU quantification. The experiments were performed using six mice per group. The skin abscess infection mouse model was revised and approved by the University Laboratory Animal Resources (ULAR) from the University of Pennsylvania (Protocol 806763).

### Deep thigh infection mouse model

Experiments were performed using six-week-old female CD-1 mice, which were rendered neutropenic by intraperitoneal application of two doses of cyclophosphamide (150 mg Kg^−1^ and 100 mg Kg^−1^) 3 and 1 days before the infection. At day 4 of the experiment, the mice were infected in their right thigh through a 100 μL intramuscular injection of *A. baumannii* ATCC19606 (in PBS at a concentration of 1×10^6^ CFU mL^−1^). The bacterial cells were grown in LB broth, washed twice with PBS solution, and diluted at the desired concentration prior to infecting the mice. The peptides were administered intraperitoneally two hours after the infection. Four-days post-infection mice were euthanized, and a uniform excision of the tissue from the right thigh was excised, homogenized using a bead beater (25 Hz for 20 min), 10-fold serially diluted, and plated on McConkey agar plates for bacterial colonies counting. The experiments were performed using six mice per group. The deep thigh infection mouse model was revised and approved by the University Laboratory Animal Resources (ULAR) from the University of Pennsylvania (Protocol 807055).

## Method details

### Encrypted peptides in archaeal proteomes

All reviewed canonical and isoform sequences of Archaea (Taxon ID: 2157) were downloaded from UniProt (https://www.uniprot.org/, access date: August 24th, 2023). We were able to obtain 19,710 protein sequences (18,677 non-redundant sequences) from 233 Archaeal organisms. Protein substrings ranging from 8-50 amino acid residues in the 18,677 sequences and containing canonical amino acids only were considered as the Archaea encrypted peptides. In total, we obtained 193,331,608 encrypted peptides from the archaeome for further studying.

### APEX 1.1

APEX is a bacterial strain-specific antimicrobial activity predictor^6^, and was trained on in-house peptide dataset and publicly available antimicrobial peptides (AMPs). Here, we updated the training data and re-trained APEX (i.e., APEX 1.1). Specifically, the updated in-house peptide dataset for training APEX contained 15,718 minimum inhibitory concentrations (MIC) values from 1,736 peptides and eleven pathogenic strains (*A. baumannii* ATCC 19606*, E. coli* ATCC 11775*, E. coli* AIC221*, E. coli* AIC222*, K. pneumoniae* ATCC 13883, *P. aeruginosa* PAO1*, P. aeruginosa* PA14*, S. aureus* ATCC 12600, methicillin-resistant *S. aureus* ATCC BAA-1556, vancomycin-resistant *E. faecalis* ATCC 700802, and vancomycin-resistant *E. faecium* ATCC 700221). Inactive data points, i.e., MIC values higher than maximum concentrations tested, were labeled as 512 μmol L^−1^. All MICs were then transformed by 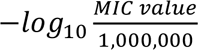. In addition to the in-house data, we curated 19,564 publicly available AMPs from DBAASP^9^, APD3^10^, and DRAMP^11^ that were not overlapped with our in-house data, as well as 9,857 non-AMPs following the instructions from Ma et al^37^, and Huang et al^38^. These publicly available AMPs and non-AMPs were used as a data augmentation strategy during APEX training^6^. We followed the original APEX paper^6^ for hyperparameter selection. The top eight APEX models were selected to create an ensemble learning where the final MIC prediction was defined as the mean predictions of the selected models.

### Physicochemical properties analysis

The six physicochemical properties of peptides, including normalized hydrophobic moment, normalized hydrophobicity, net charge, disordered conformation propensity, propensity to aggregation *in vitro*, and amphiphilicity index, were obtained from the DBAASP server^9^. Note that Eisenberg and Weiss scale^39^ was chosen as the hydrophobicity scale.

### Phylogenetic tree visualization and phylogenetic distance

To obtain the phylogenetic tree, the taxon IDs of 233 Archaeal organisms obtained from UniProt were uploaded to NCBI Taxonomy Common Tree (https://www.ncbi.nlm.nih.gov/Taxonomy/CommonTree/wwwcmt.cgi). Of note, 201 out of 233 Archaea organisms were successfully retrieved. The resulted tree file from NCBI was then visualized via iTOL (https://itol.embl.de/). The distance of two nodes in the phylogenetic tree is defined as the length of the shortest path between these two nodes.

### Peptide sequence similarity

Let *SW*(*i*, *j*) denote the Smith-Waterman alignment score^40^ between two protein sequences *i* and *j*. We define the peptide sequence similarity between *i* and *j* as 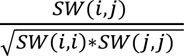.

### Peptide sequence space visualization

Given a peptide dataset, we calculated a similarity matrix to represent the pairwise sequence similarities among the peptides. We then applied Uniform Manifold Approximation and Projection (UMAP) to transform this similarity matrix into a two-dimensional space. This transformed space serves as a proxy for the peptide sequence space, allowing us to visualize the distribution of peptides within it.

### Archaeasins selection

APEX 1.1 was used to predict the antimicrobial activity for the 193,331,608 encrypted peptides derived from the archaeome. We used the mean MIC value against the eleven pathogen strains to rank and select the encrypted peptides for chemical synthesis and experimental validation (**Data S2**). When selecting the peptides, we also make sure they met the following criteria:

1. The selected peptide should have <70% sequence similarity to all in-house peptides and publicly available AMPs.
2. The selected peptides themselves should have <70% sequence similarity.
3. The selected peptide should have ≤64 μmol L^−1^ mean MIC by prediction.

### Peptide Synthesis

All peptides used in the experiments were purchased from AAPPTec and synthesized by solid-phase peptide synthesis using the Fmoc strategy.

### Minimal inhibitory concentration determination

Broth microdilution assays were performed to determine the minimum inhibitory concentration (MIC) values of each peptide. Peptides were added to nontreated polystyrene microtiter 96-well plates and 2-fold serially diluted in sterile water from 1 to 64 μmol L^−1^. Bacterial inoculum at 4×10^6^ CFU mL^−1^ in LB or BHI medium was mixed 1:1 with the peptide. The MIC was defined as the lowest concentration of peptide able to completely inhibit the bacterial growth after 24 h of incubation at 37 °C. All assays were done in three independent replicates.

### Circular dichroism experiments

The circular dichroism experiments were conducted using a J1500 circular dichroism spectropolarimeter (Jasco) in the Biological Chemistry Resource Center (BCRC) at the University of Pennsylvania. Experiments were performed at 25 °C, the spectra graphed are an average of three accumulations obtained with a quartz cuvette with an optical path length of 1.0 mm, ranging from 260 to 190 nm at a rate of 50 nm min^−1^ and a bandwidth of 0.5 nm. The concentration of all EPs tested was 50 μmol L^−1^, and the measurements were performed in water, a mixture of trifluoroethanol (TFE) and water in a 3:2 ratio, and sodium dodecyl sulfate (SDS) in water at 10 mmol L^−1^, with respective baselines recorded prior to measurement. A Fourier transform filter was applied to minimize background effects. Secondary structure fraction values were calculated using the single spectra analysis tool on the server BeStSel^16^. Ternary plots were created in https://www.ternaryplot.com/ and subsequently edited.

### Synergy assays

Combinations of two encrypted peptides from the same protein were tested against *A. baumannii* ATCC 19606 strains using the checkboard assay. Briefly, two-fold serial dilutions of each peptide were orthogonally mixed and incubated with a bacterial suspension at a final concentration of 2×10^6^ CFU mL^−1^ in LB for 24 h at 37 °C. The fractional inhibitory concentration indexes (FICI) were defined to attribute whether the interactions between peptides were synergistic (FICI≤0.5), additive (0.5>FICI≥1), or indifferent (FICI>1).

### Outer membrane permeabilization assays

N-phenyl-1-napthylamine (NPN) uptake assay was used to evaluate the ability of the peptides to permeabilize the bacterial outer membrane. Inocula of *A. baumannii* ATCC 19606 were grown to an OD at 600 nm of 0.4 mL^−1^, centrifuged (10,000 rpm at 4 °C for 10 min), washed and resuspended in 5 mmol L^−1^ HEPES buffer (pH 7.4) containing 5 mmol L^−1^ glucose. The bacterial solution was added to a white 96-well plate (100 μL per well) together with 4 μL of NPN at 0.5 mmol L^−1^. Consequently, peptides diluted in water were added to each well, and the fluorescence was measured at λex = 350 nm and λem = 420 nm over time for 45 min. The relative fluorescence was calculated using the untreated control (buffer + bacteria + fluorescent dye) and polymyxin B (positive control) as baselines and the following equation was applied to reflect % of difference between the baselines and the sample:

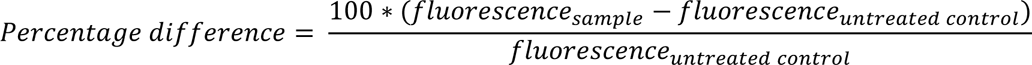

### Cytoplasmic membrane depolarization assays

The cytoplasmic membrane depolarization assay was performed using the membrane potential-sensitive dye 3,3’-dipropylthiadicarbocyanine iodide (DiSC3-5). *A. baumannii* ATCC 19606 and *P. aeruginosa* PAO1 in the mid-logarithmic phase were washed and resuspended at 0.05 OD mL^−1^ (optical value at 600 nm) in HEPES buffer (pH 7.2) containing 20 mmol L^−1^ glucose and 0.1 mol L^−1^ KCl. DiSC3-5 at 20 μmol L^−1^ was added to the bacterial suspension (100 μL per well) for 15 min to stabilize the fluorescence which indicates the incorporation of the dye into the bacterial membrane, and then the peptides were mixed 1:1 with the bacteria to a final concentration corresponding to their MIC100 values. Membrane depolarization was then followed by reading changes in the fluorescence (λex = 622 nm, λem = 670 nm) over time for 60 min. The relative fluorescence was calculated using the untreated control (buffer + bacteria + fluorescent dye) and polymyxin B (positive control) as baselines and the following equation was applied to reflect % of difference between the baselines and the sample:

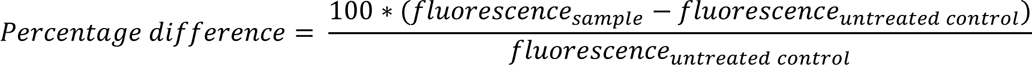

### Hemolytic activity assays

To evaluate the release of hemoglobin from human erythrocytes upon treatment of each of the encrypted peptides, human red blood cells (RBCs) were obtained from ZenBio (male donor, blood type A^−^) obtained from heparin anti-coagulated blood. RBCs were washed with PBS (pH 7.4) four times by centrifugation at 800 g for 10 min. Aliquots of 200-fold diluted cells (75 μL) were mixed with peptide solution (0.78-100 μmol L^−1^; 75 μL), and the mixture was incubated for 4 h at room temperature. After the incubation, the plate was centrifuged at 1,300 ×g for 10 min to precipitate cells and debris, and 100 μL of supernatant from each well were transferred to a new 96-well plate for absorbance reading (405 nm) using an automatic plate reader. The percentage of hemolysis was defined by comparison with negative control (samples containing PBS) and positive control [samples containing 1% (v/v) SDS in PBS solution].

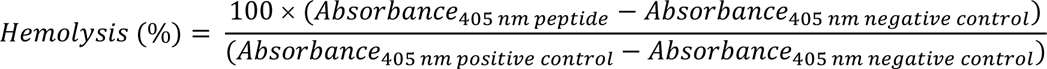

### Quantification and statistical analysis

#### Reproducibility of the experimental assays

All assays were performed in three independent biological replicates as indicated in each figure legend and in the Experimental Models and Methods details sections. The values obtained for hemolytic activity were estimated by non-linear regression based on the screen of peptides in a gradient of concentrations and represent the hemolytic concentration values needed to lyse and kill 50% of the cells present in the experiment. In the skin abscess and thigh infection mouse models, we used six mice per group following established protocols approved by the University Laboratory of Animal Resources (ULAR) of the University of Pennsylvania.

#### Statistical tests

In the mouse experiments, all the raw data were log10 transformed and the statistical significance was determined using one-way ANOVA followed by Dunnett’s test. All the p values are shown for each of the groups, all groups were compared to the untreated control group.

#### Statistical analysis

All calculation and statistical analyses of the experimental data were conducted using GraphPad Prism v.10.3. Statistical significance between different groups was calculated using the tests indicated in each figure legend. No statistical methods were used to predetermine sample size.

## Supplemental figures and tables

**Supplementary Table S1.** Pearson correlation between predicted and experimentally-determined MICs by APEX and APEX 1.1 on the 80 Archaea encrypted peptides we synthesized and validated.

| Bacterial strains | APEX1.1 | APEX |
| --- | --- | --- |
| *A. baumannii* ATCC 19606 | 0.3951 | 0.2038 |
| *E. coli* ATCC 11775 | 0.2488 | 0.1201 |
| *E. coli* AIC221 | 0.2553 | 0.0760 |
| *E. coli* AIC222 | 0.1886 | 0.1253 |
| *K. pneumoniae* ATCC 13883 | 0.1020 | -0.0394 |
| *P. aeruginosa* PAO1 | 0.3037 | 0.3657 |
| *P. aeruginosa* PA14 | 0.4662 | 0.3608 |
| *S. aureus* ATCC 12600 | 0.3213 | 0.1622 |
| Methicillin-resistant *S. aureus* ATCC BAA-1556 | 0.1029 | 0.0698 |
| Vancomycin-resistant *E. faecalis* ATCC 700802 | 0.1772 | 0.0824 |
| Vancomycin-resistant *E. faecium* ATCC 700221 | 0.3634 | 0.2960 |
| Macro average | 0.2659 | 0.1657 |
| Micro average | 0.5033 | 0.4455 |

**Supplementary Table S2.** Spearman correlation between predicted and experimentally-determined MICs by APEX and APEX 1.1 on the 80 Archaea encrypted peptides we synthesized and validated.

| Bacterial strains | APEX1.1 | APEX |
| --- | --- | --- |
| *A. baumannii* ATCC 19606 | 0.3971 | 0.2142 |
| *E. coli* ATCC 11775 | 0.2282 | 0.1328 |
| *E. coli* AIC221 | 0.2405 | 0.0707 |
| *E. coli* AIC222 | 0.1944 | 0.0935 |
| *K. pneumoniae* ATCC 13883 | 0.1314 | -0.0559 |
| *P. aeruginosa* PAO1 | 0.2882 | 0.3380 |
| *P. aeruginosa* PA14 | 0.4932 | 0.3906 |
| *S. aureus* ATCC 12600 | 0.2575 | 0.1442 |
| Methicillin-resistant *S. aureus* ATCC BAA-1556 | 0.0122 | 0.0544 |
| Vancomycin-resistant *E. faecalis* ATCC 700802 | 0.1785 | 0.1539 |
| Vancomycin-resistant *E. faecium* ATCC 700221 | 0.3242 | 0.2330 |
| Macro average | 0.2496 | 0.1609 |
| Micro average | 0.4908 | 0.4340 |

**Figure S1.**
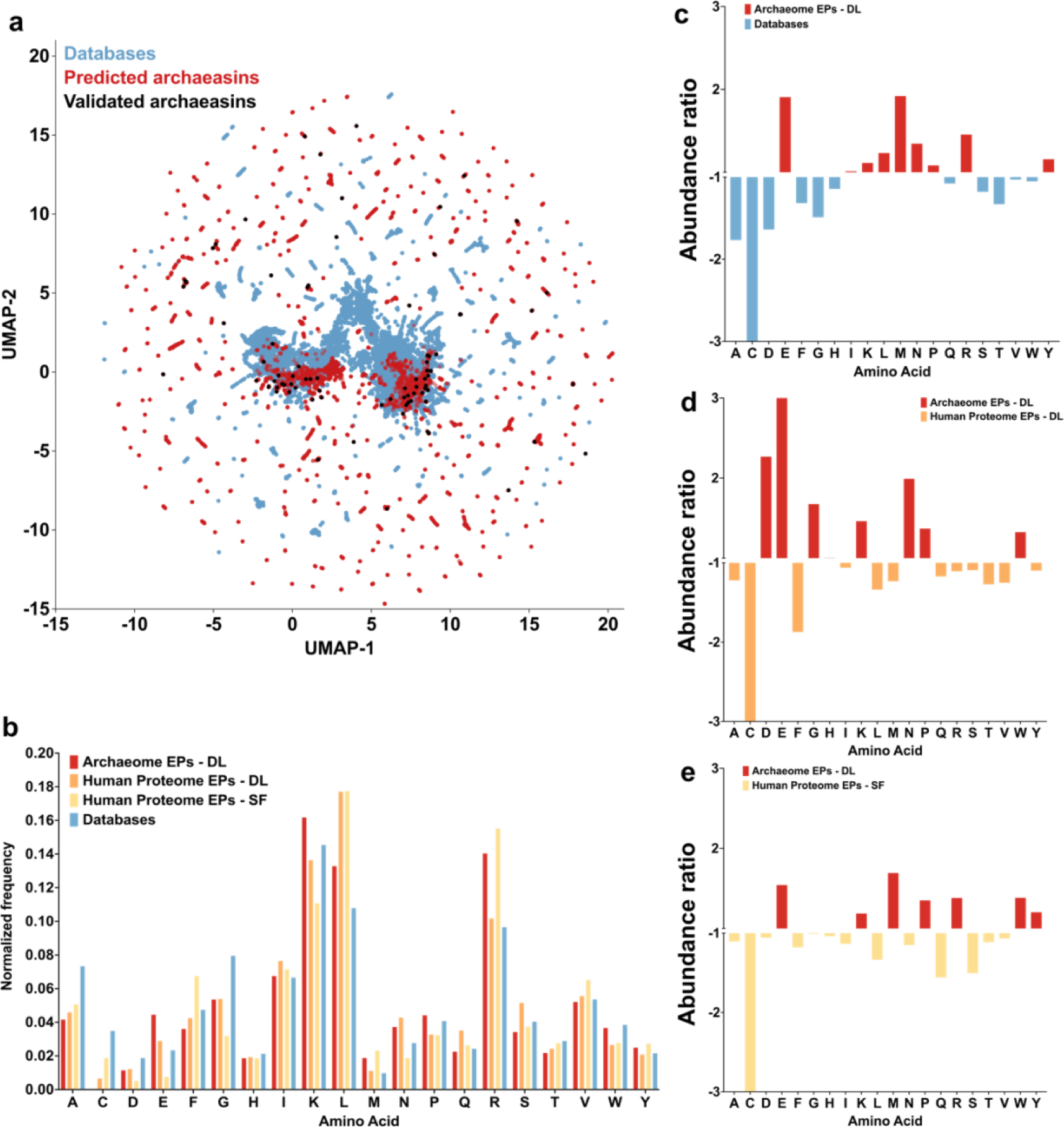
Sequence features of the archaeasins. **(a)** Bidimensional sequence space visualization of peptide sequences found in DBAASP, all predicted antimicrobial EPs discovered by APEX, and peptides that were synthesized and validated in this study. Uniform Manifold Approximation and Projection (UMAP) was used to reduce the feature representation to two dimensions for visualization purposes. **(b)** Amino acid frequency in archaeasins compared with known AMPs from DBAASP, APD3, and DRAMP 3.0 databases, and other encrypted peptides from the human proteome discovered by APEX^6^ and a scoring function^4^. **(c-e)** Relative abundance of the amino acid content of Archaea encrypted peptides identified by APEX (red) and **(c)** AMPs from the DBAASP, APD3, and DRAMP 3.0 database (blue), **(d)** human proteome encrypted peptides discovered by APEX (orange), and **(e)** human encrypted peptides found by the scoring function (yellow). The frequency of amino acid was normalized by the total number of amino acid residue counts.

**Figure S2.**
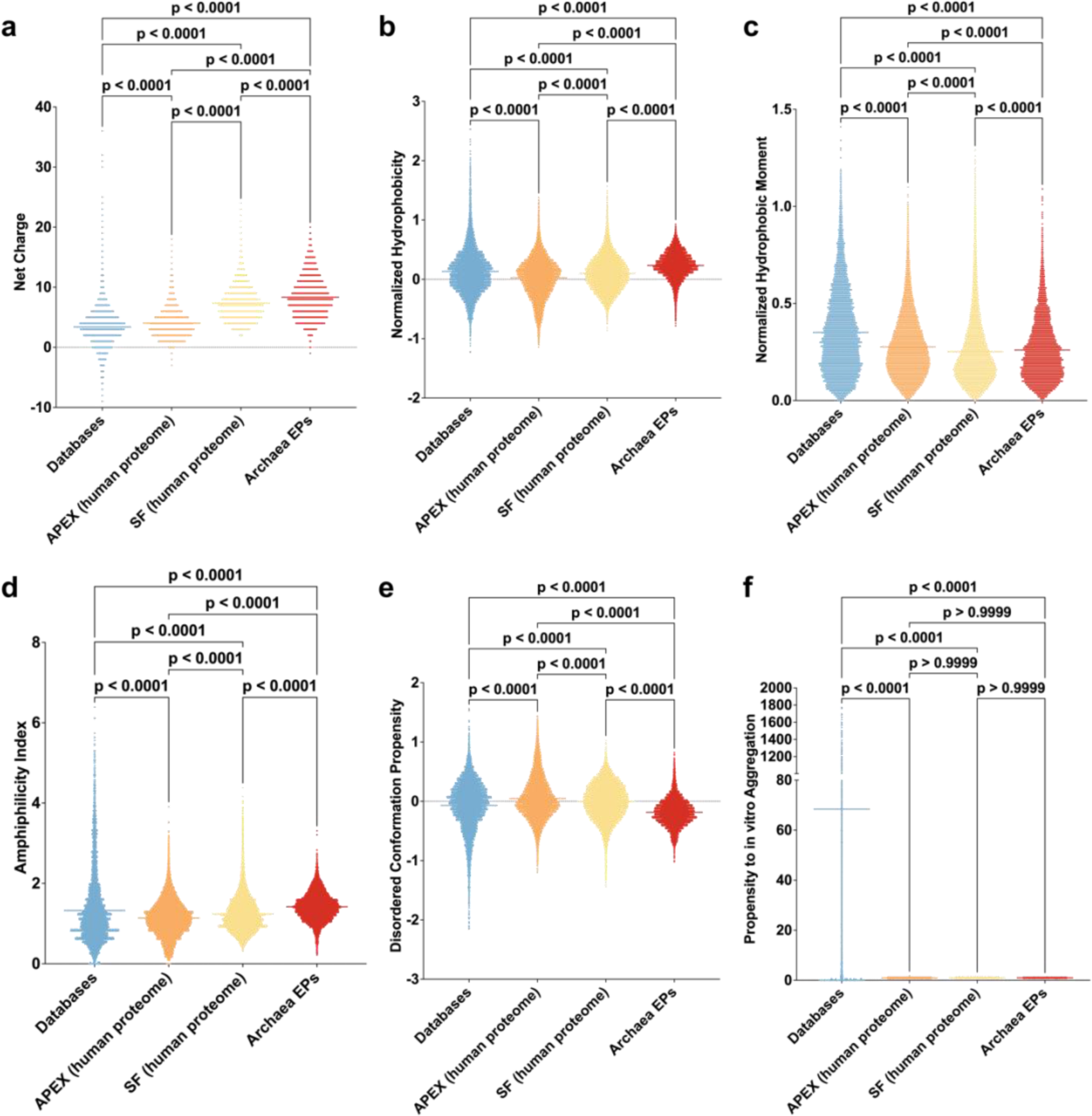
Physicochemical features of archaeasins compared to AMPs from databases (DBAASP, APD3, and DRAMP 3.0), and encrypted peptides from human proteins identified by APEX and a scoring function. **(a)** Net charge and **(b)** normalized hydrophobicity. **(c)** Hydrophobic moment normalized by peptide length, reflecting the amphipathicity of the molecules, which directly influences their interactions with bacterial membranes. **(d)** Amphiphilicity index and **(e)** disordered conformation propensity, both of which are closely correlated with the mechanism of action, specifically how peptides interact with membrane lipids to exert antimicrobial activity. **(f)** Propensity to aggregate *in vitro*, correlated with the supramolecular arrangement of the molecules and potential toxicity. Statistical significance was determined using two-tailed t-tests followed by the Mann-Whitney test; p values are shown in the graph. The solid line within each box represents the mean value for each group.

**Figure S3.**
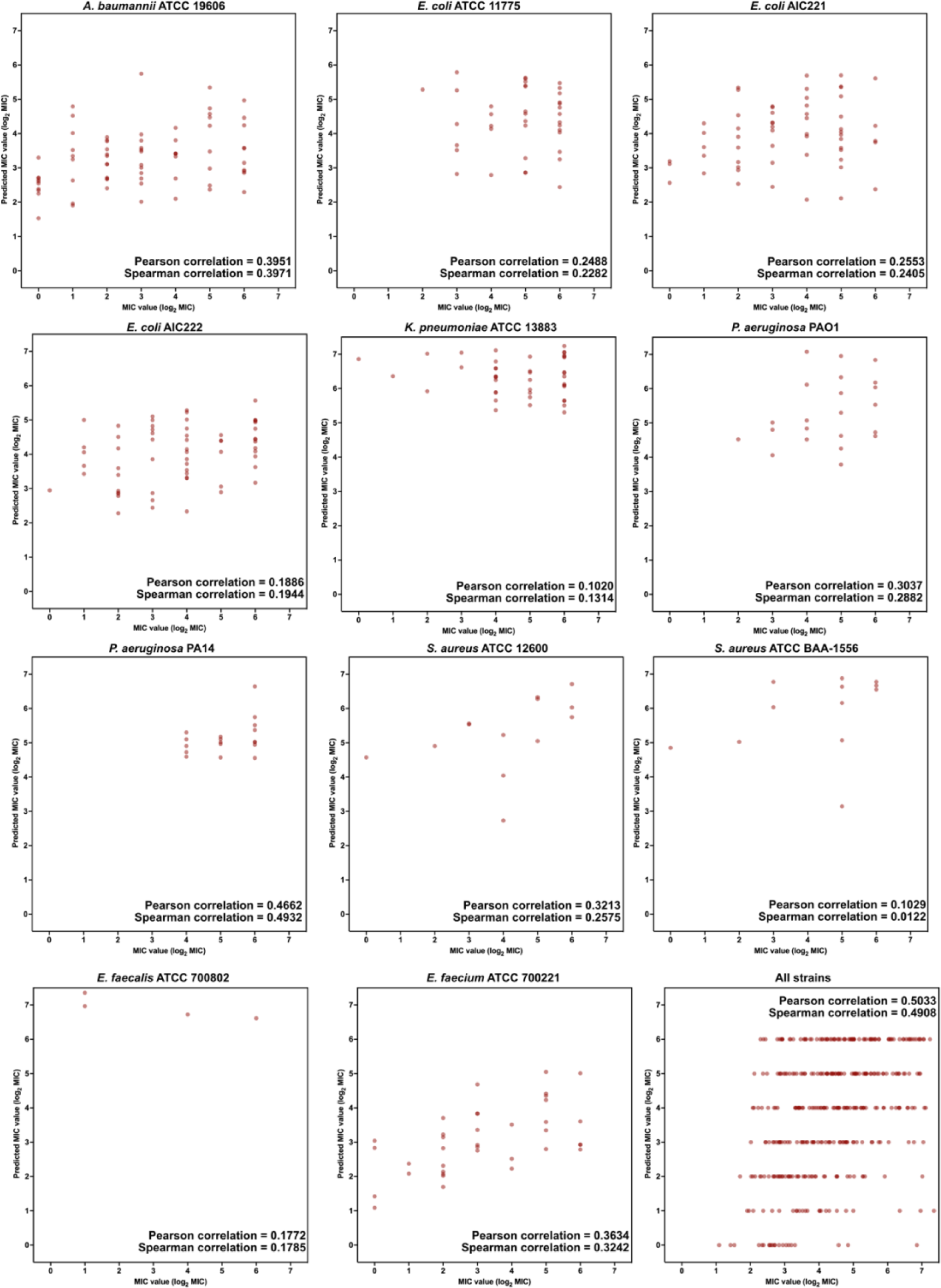
Predicted vs. experimental MIC values of the archaeasins identified by APEX against various pathogens. Each peptide is represented by a red circle on the scatter plot. Pearson and Spearman correlation values are provided as insets.

**Figure S4.**
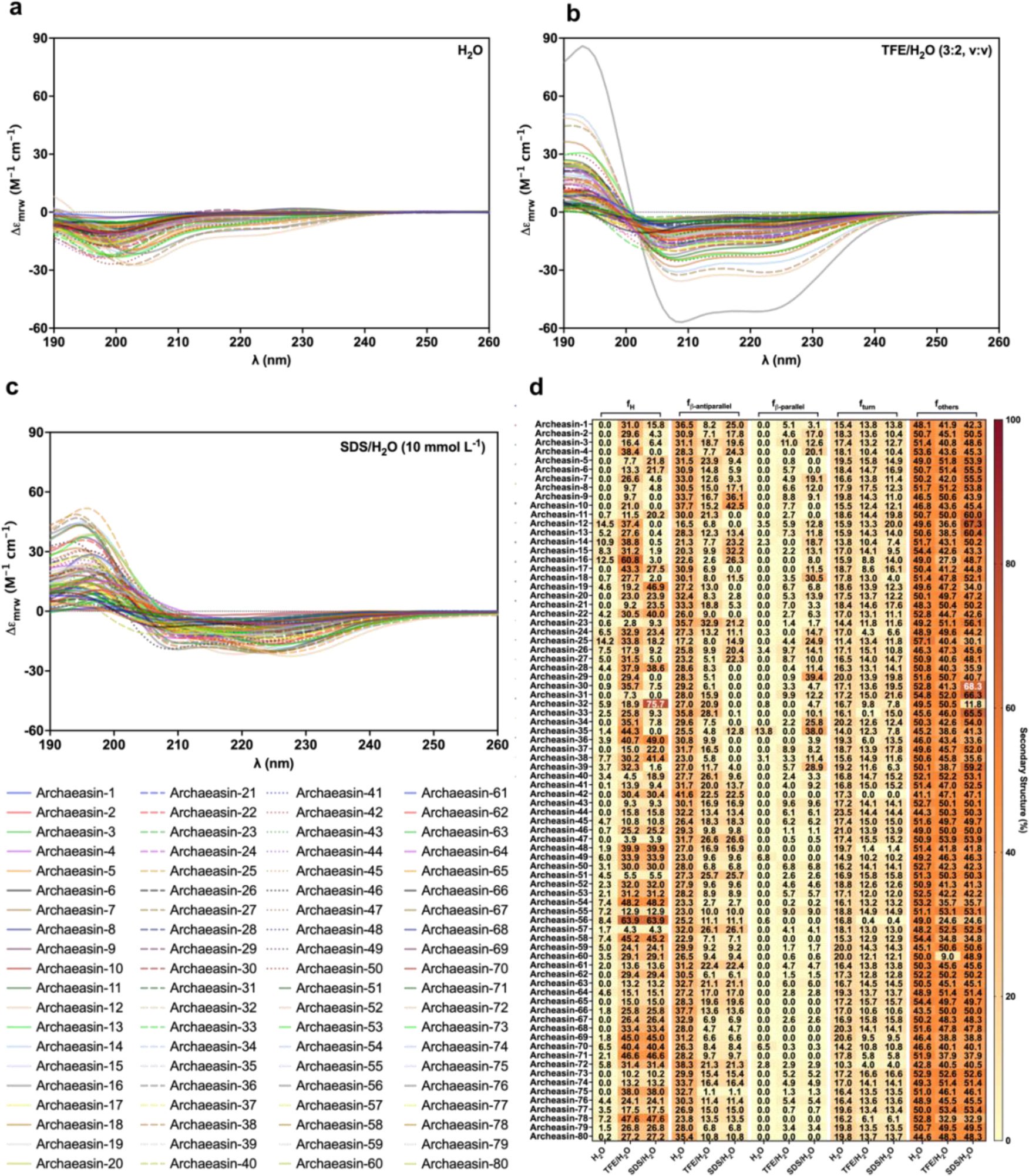
Circular dichroism spectra of archaeasins. Circular dichroism experiments were conducted with encrypted peptides from the archaeome using a J-1500 Jasco circular dichroism spectrophotometer. The spectra were recorded in three different media: **(a)** water, **(b)** 60% trifluoroethanol in water, and **(c)** sodium dodecyl sulfate (SDS) in water (10 mmol L^−1^), after three accumulations at 25 °C, using a 1mm path length quartz cell, between 260 and 190 nm at 50 nm min^−1^, with a bandwidth of 0.5 nm. The concentration of all peptides tested was 50 μmol L^−1^. **(d)** Heatmap with the percentage of secondary structure found for each peptide in three different solvents: water, 60% trifluoroethanol (TFE) in water, and SDS (10 mmol L^−1^) in water. Secondary structure fraction was calculated using the BeStSel server^16^.

**Figure S5.**
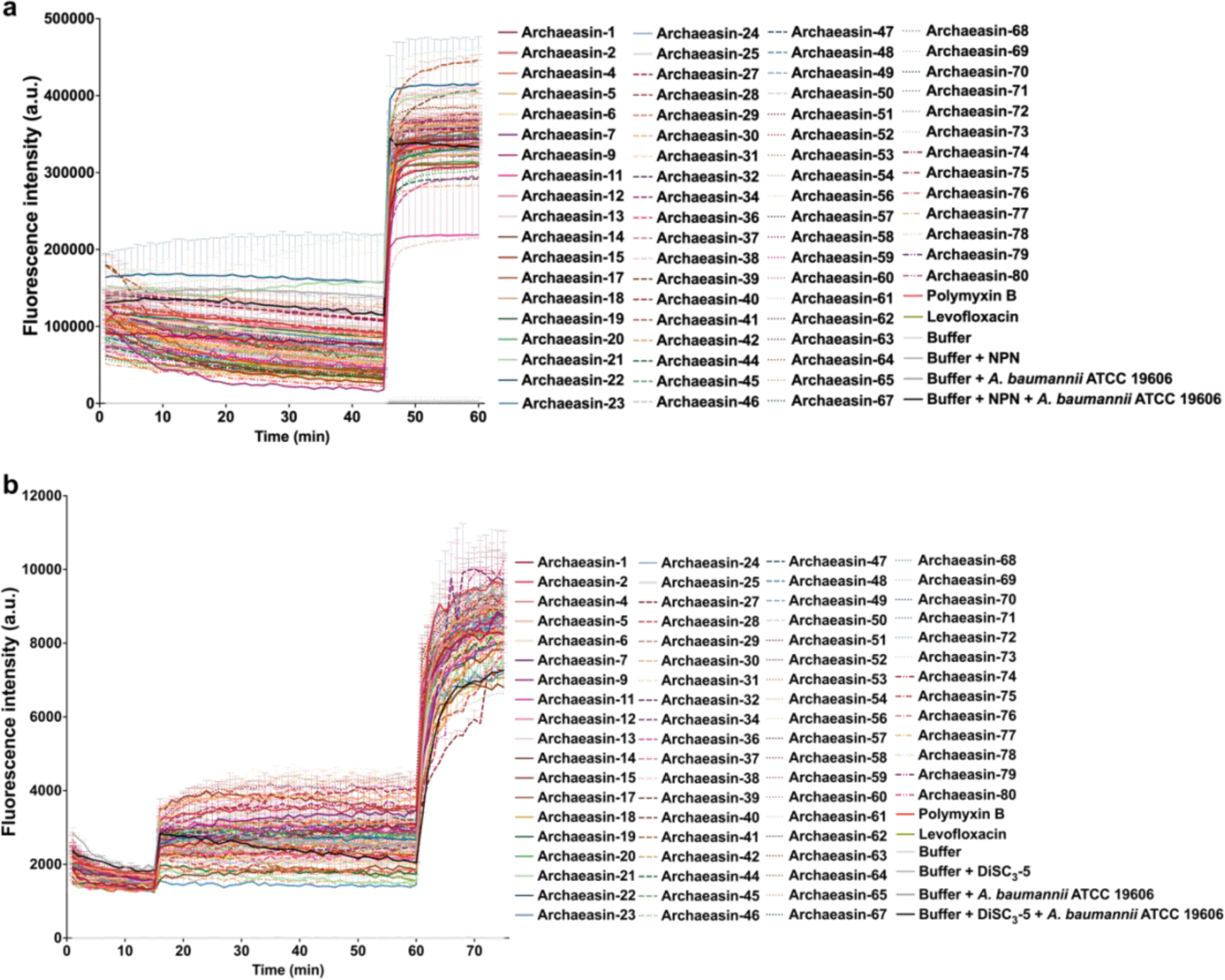
Outer membrane permeabilization and cytoplasmic membrane depolarization of *A. baumannii* ATCC 19606 induced by archaeasins. **(a)** Outer membrane permeabilization was assessed using the probe 1-(N-phenylamino)naphthalene (NPN), showing the permeabilization effects of Archaea-encoded encrypted peptides active against *A. baumannii* ATCC 19606. **(b)** Membrane depolarization assays were performed using the hydrophobic probe 3,3′-dipropylthiadicarbocyanine iodide [DiSC3-(5)] on all archaeasins active against *A. baumannii* ATCC 19606. Polymyxin B served as a positive control, while buffer, buffer with the probe, and buffer with both probe and bacteria were used as baseline controls for fluorescence. The panels display the raw fluorescence intensity data obtained from the experiments. Error bars are the standard deviation obtained from the three replicates.

**Figure S6.**
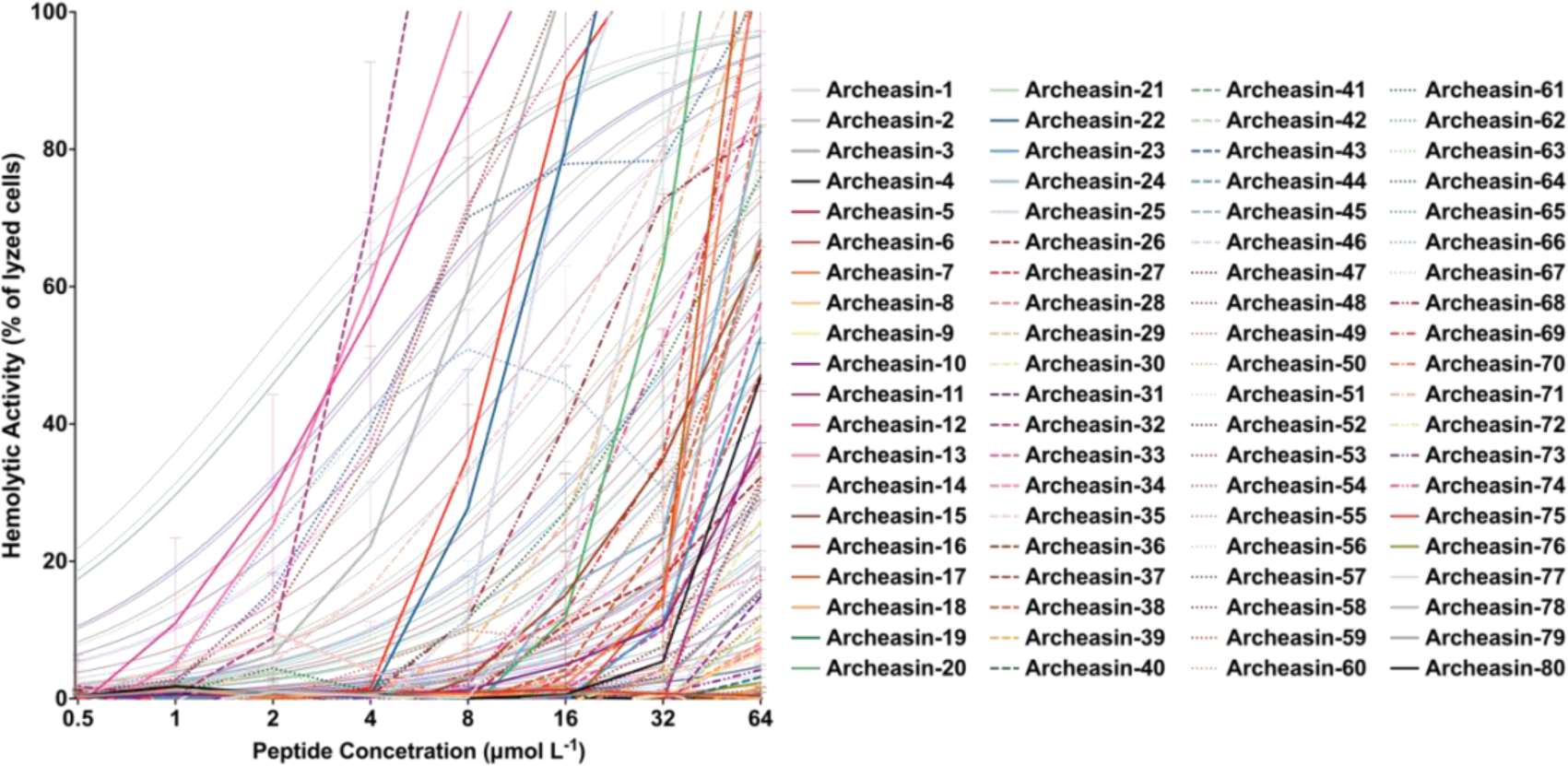
Hemolytic activity of archaeasins. Hemolytic activity was assessed by exposing red blood cells to archaeasins, with lysis determined by absorbance readings after 4 hours. The peptides were added to plates and subjected to a two-fold dilution series (ranging from 64 to 4 μmol L^−1^) before exposure to the red blood cells.

**Figure S7.**
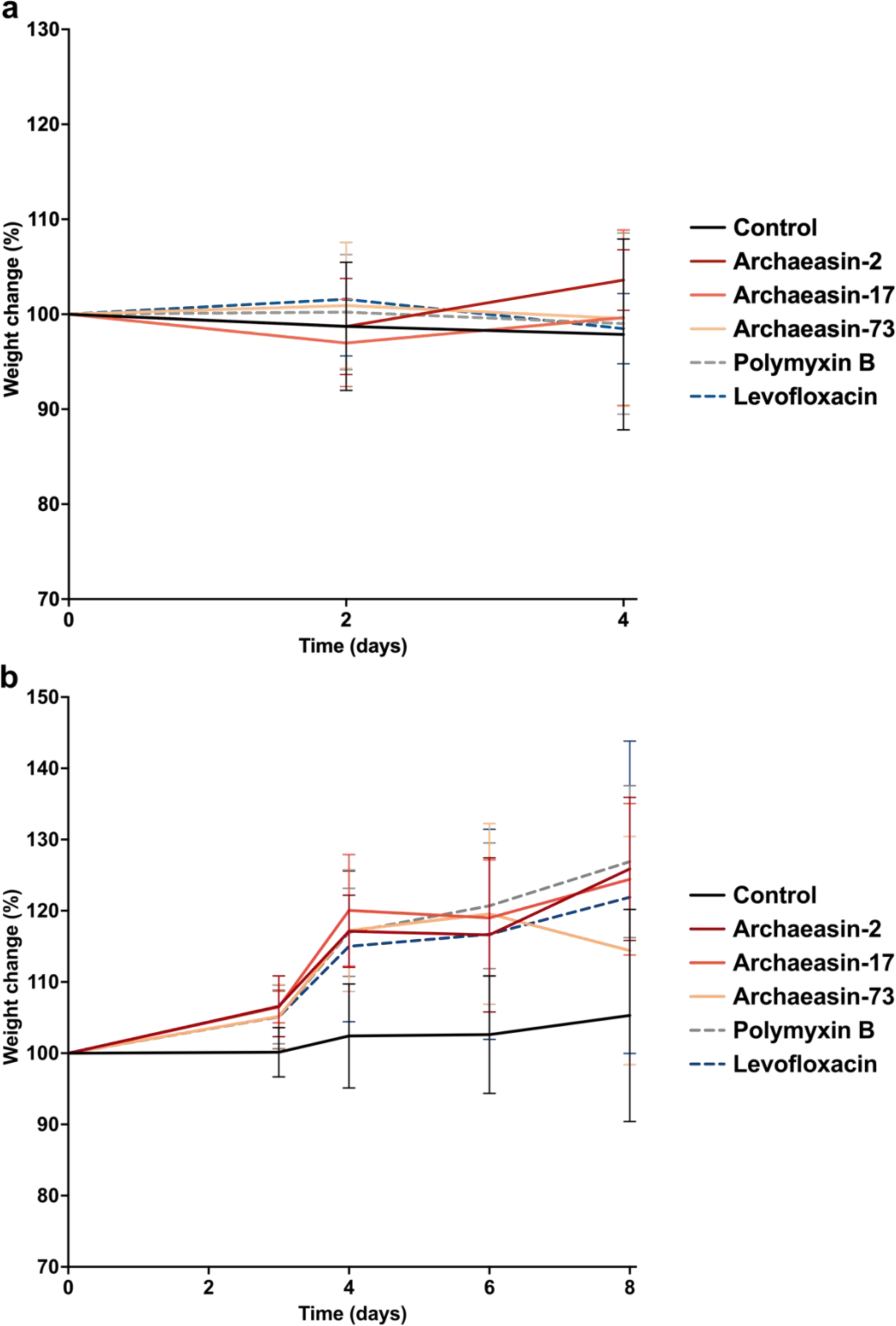
Weight change monitoring in both skin abscess and deep thigh infection mouse models infected with *A. baumannii*. Mouse weight was monitored throughout the duration of the **(a)** skin abscess model (4 days total) and the **(b)** deep thigh infection model (8 days total) to assess potential toxic effects of both the bacterial load and the archaeasins.

